# Pervasive prophage recombination occurs during evolution of spore-forming *Bacilli*

**DOI:** 10.1101/2020.05.06.055103

**Authors:** Anna Dragoš, Priyadarshini B., Zahraa Hasan, Mikael Lenz-Strube, Paul J Kempen, Gergely Maróti, Charlotte Kaspar, Baundauna Bose, Briana M. Burton, Ilka B Bischofs, Ákos T. Kovács

**Author notes:** Corresponding authors: Bacterial Interactions and Evolution Group, Department of Biotechnology and Biomedicine, Technical University of Denmark, Søltofts Plads Building 221, 2800 Kongens Lyngby, Denmark.

## Abstract

Phages are the main source of within-species bacterial diversity and drivers of horizontal gene transfer, but we know little about the mechanisms that drive genetic diversity of these mobile genetic elements (MGEs). Recently, we showed that a sporulation selection regime promotes evolutionary changes within SPβ prophage of *Bacillus subtilis*, leading to direct antagonistic interactions within the population. Herein, we reveal that under a sporulation selection regime, SPβ recombines with low copy number phi3Ts phage DNA present within the *B. subtilis* population. Recombination results in a new prophage occupying a different integration site, as well as the spontaneous release of virulent phage hybrids. Analysis of *Bacillus* sp. strains suggests that SPβ and phi3T belong to a distinct cluster of unusually large phages inserted into sporulation-related genes that are equipped with a spore-related genetic arsenal. Comparison of *Bacillus* sp. genomes indicates that similar diversification of SPβ-like phages takes place in nature. Our work is a stepping stone toward empirical studies on phage evolution, and understanding the eco-evolutionary relationships between bacteria and their phages. By capturing the first steps of new phage evolution, we reveal striking relationship between survival strategy of bacteria and evolution of their phages.

## Introduction

Bacteriophages are major regulators of bacteria. Each day, approximately half of bacterial biomass is killed by lytic phages, imposing a constant predator-prey evolutionary arms race [1–3]. Phages reside in 40-50% of bacterial genomes as prophage elements [4], serving as a main source of intra-species genetic diversity and gene transfer agents [5]. Despite their abundance and relevance, we still understand very little about ecological and evolutionary imprint of prophages. This knowledge gap is particularly paramount for non-pathogenic bacteria, such as widely applied biocontrol and probiotic strains, which often (e.g. *Bacillus* genus) can be extremely rich in prophages [6]. Moreover, between-strains differences in prophage elements in certain *Bacillus* species (e.g. *Bacillus subtilis*) leads to social ‘incompatibility’, which manifests in strong competitive interactions and physical barriers between bacterial swarms [7, 8]. Currently, we do not understand what promotes prophage diversity within closely related strains.

It is believed that phages predominantly evolve through recombination [9–11], but the rate of horizontal gene transfer (HGT) can vary substantially, depending on phage lifestyle of host phylum [12]. Gene gain/loss by HGT seems to be more common in Firmicutes or Bacteroides phages, compared to Cyanophages, and it is more pronounced in phages with temperate lifestyle [12–14]. Recombination between phages or between phages and their hosts can be either homologous [15, 16] or non-homologous [17], relying on phage- or host-encoded recombinases [15, 18]. It was proposed that gene shuffling regularly occurs between (functional or defective) prophages and phages that co-infect the same host bacterium [15]. Prophages of naturally competent bacteria (e.g. *B. subtilis*) can recombine with foreign phage DNA via transformation [19]. Finally, phages can randomly or specifically incorporate fragments of host chromosomes via generalised [20, 21] or specialised [22, 23] transduction, respectively, thereby contributing to the spread of antibiotic resistance [21] or virulence genes [24]. Although evidence from comparative genomics indicates frequent phage recombination [25-27], there is little experimental research in this topic. Phage recombination has been experimentally studied using few models, predominantly *Salmonella typhimurium* P22 with *Escherichia coli* lambdoid phages [28, 29], or *Lactococcus lactis* phages sampled across decades from diary production line [25]. Therefore, empirical research on prophage evolution is limited.

Interestingly, certain prophages of *Bacilli* undergo genetic rearrangements upon host development, acting as so-called phage regulatory switches (RSs) [30]. RS can switch between integrated and extrachromosomal forms to modulate reproduction and survival of their hosts [30]. Certain *B. subtilis* strains carry an SPβ prophage or its derivative that integrates into an *spsM* gene, encoding for polysaccharide synthase [31, 32]. Immediately prior to sporulation the prophage undergoes precise excision and circularisation, allowing *spsM* reconstitution and expression in the sporulating mother cell. The resulting *spsM*-related polysaccharide eventually becomes part of the spore coat, contributing to spore dispersability [31]. Besides SPβ, another prophage-like element named *skin* also undergoes excision in the mother cell, allowing reconstitution of *sigK* encoding a late sporulation sigma factor that is necessary for completing the sporulation process [33-35]. Similar mother cell-specific excisions have been observed in other *Bacillus* sp. [31] and in *Clostridium* sp. [36, 37], but we do not understand what drives such a distinctive relationship between spore-forming hosts and their phages, nor what eco-evolutionary consequences this has. Interestingly, SPβ and *skin* both encompass genes relevant for sporulation, including *sspC* that is crucial for spore DNA protection and repair [38, 39], and the *rapE-phrE* signalling system involved in sporulation initiation [40, 41]. Furthermore, certain prophages can improve or even restore sporulation in *B. subtilis* [42], suggesting the possibility of a cooperative relationship between certain phages and spore-formers.

We recently demonstrated that under a repeated imposed sporulation selection regime, SPβ prophages of *B. subtilis* undergo major genetic rearrangements, giving rise to new hybrid phages [43]. Normally, the lytic cycle of SPβ prophages is blocked by the ICEBs1 (Integrative and Conjugative Element of *B. subtilis*) conjugative element [44], but in the evolved strains, phages are released spontaneously, killing or lysogenizing the original host [43].

Herein, we investigated the triggering cause of diversification and spontaneous release of SPβ prophages and sought evidence for similar diversification of SPβ taking place in nature. Using experimental evolution, *de novo* genome sequencing and testing, we showed that barely detectable, low copy number phage DNA residing in certain *B. subtilis* strains can propagate under an appropriate selection regime and recombine with indigenous prophages. The new prophages modulate host development, most likely through regulatory genes. Bioinformatic comparison of available *Bacillus* sp. genomes demonstrated that similar recombination may frequently occur in nature between SPβ and related phages. Our work shows how diversification of prophages through recombination can drive early diversification of bacterial populations.

## Materials and Methods

### Strains and cultivation conditions

Supplementary Table S3 describes the bacterial strains used in this study and Supplementary Table S4 lists all phages used in this work. Plasmids and oligonucleotides used for cloning purposes to construct some of the strains used here are listed in Supplementary Table S5. Strains were routinely maintained in lysogeny broth (LB) medium (LB-Lennox, Carl Roth; 10 g/l tryptone, 5 g/l yeast extract, and 5 g/l NaCl).

Strain DTUB200 was obtained by lysogenization of DK1042 (WT NCBI 3610) with a phage phi3T obtained from CU1065. DTUB201 (ΔSPβ) was obtained by transforming DK1042 with gDNA obtained from SPmini and selecting for erythromycin-resistant colonies. Strain DTUB202 (Δ*kamA*) was obtained by transforming DK1042 with gDNA obtained from BKK19690 and selecting for kanamycin-resistant colonies. Strains DTUB251 and DTUB254 were obtained by transforming DK1042 with plasmids pDR111_rapX and pDR111_spsX, respectively. Plasmid pDR111 was linearized using oAD47/pAD48 primers, *rapX* and *spsX* were amplified from B410mB using oAD49/oAD50 and pAD71/oAD72 primers, respectively (see Suppl. table 5). Next, the linearized pDR111 was ligated with *rapX* and with *spsX* using Gibson Assembly® Master Mix (New England BioLabs), resulting in pDR111_rapX and pDR111_spsX. All modifications of DK1042 were verified by PCR followed by Sanger sequencing.

### Genome sequencing and analysis

Phage sequencing was performed by Illumina MiSeq instrument and a 2×250 nt paired-end chemistry (MiSeq Reagent Kit v2 (500-cycles). Primary data analysis (base-calling) was carried out with Bbcl2fastq^ software (v2.17.1.14, Illumina). In vitro fragment libraries were prepared using the NEBNext® Ultra™ II DNA Library Prep Kit for Illumina. Reads were quality and length trimmed in CLC Genomics Workbench Tool 11.0 and *de novo* genome assembly was performed using SPAdes-3.13.0-Linux and CLC Genomics Workbench 11.0.

*De novo* sequencing and assembly of B310mA, B410mB and B410wtB genomes was performed by Functional Genomics Center Zurich, from genomic DNA of exponentially grown cultures, extracted using the EURex Bacterial and Yeast Genomic DNA Kit. Sequencing of 168 ancestor (ancestor of B310mA, B410mB and B410wtB) was described in our previous manuscript [43].

Evolved PY79 strains (presented in Suppl. dataset 2) were obtained as previously described [45]. Samples for whole-genome sequencing were prepared according to the Illumina Multiplexing Sample Preparation Guide, using NEBNext reagents and Illumina’s indexed primers. Sequencing was performed by the Bauer Core Facility at Harvard University. Mapping of raw fastq reads to reference PY79 genome (NC_022898.1) was performed using Bowtie2 [46, 47]. The alignment was sorted using SAMtools [48, 49], data filtering and SNP variant calling was performed using the bcftools package. Mapping of raw fastq reads to phi3T genome (KY030782.1) was performed using Bowtie2 in Galaxy platform (https://cpt.tamu.edu/galaxy-pub) and coverage was visualized in the browser using Trackster tool. Mapping of raw SOLiD sequencing reads (168_anc_) to unique phi3Ts fragments was performed using CLC Genomics Workbench 11.0.1. Short phi3T fragments, to which fastq could be mapped, showed over 90% sequence identity to PY79 genome, as confirmed by BLAST. All bacterial and phage genomes sequenced during this work, where deposited at NCBI database as completed genomes and/or raw sequencing data (Table 1).

**Table 1.** List of bacterial strain and phages subjected to genome sequencing with corresponding NCBI accession numbers.

| Name of bacterial strain/phage | Data | Accession number |
| --- | --- | --- |
| B310mA | Complete genome | CP051860 |
| B410mB | Complete genome | CP053102* |
| B410wtB | Complete genome | CP052842* |
| B310mA | Sequencing reads (Illumina) | SRR11561554 |
| B410mB | Sequencing reads (Illumina) | SRR1156151 |
| B410wtB | Sequencing reads (Illumina) | SRR11561552 |
| 168<sub>ancestor</sub> | Sequencing reads (SOLiD) | SRR11559011 |
| NCBI 3610 | Sequencing reads (Illumina) | SRR11559035 |
| 15.1 | Sequencing reads (Illumina) | SRR11566357 |
| 16.1 | Sequencing reads (Illumina) | SRR11566355 |
| 16.2 | Sequencing reads (Illumina) | SRR11566354 |
| phi3Ts | Complete genome | MT366945 |
| Hyb1<sup>phi3Ts-SPβ</sup> | Complete genome | MT366946 |
| Hyb2<sup>phi3Ts-SPβ</sup> | Complete genome | MT366947 |
| Hyb3<sup>phi3Ts-SPβ</sup> | Complete genome | MT366948 |
| phi3Ts | Sequencing reads (Illumina) | SRR11587866 |
| Hyb1<sup>phi3Ts-SPβ</sup> | Sequencing reads (Illumina) | SRR11587864 |
| Hyb2<sup>phi3Ts-SPβ</sup> | Sequencing reads (Illumina) | SRR11587865 |
| Hyb3<sup>phi3Ts-SPβ</sup> | Sequencing reads (Illumina) | SRR11587867 |

\*extrachromosomal phage fragments: B410wtB - CP052843; B410mB – supplementary dataset 4.

Raw sequencing data of PY79 strains [45] and available from B. Burton. Raw sequencing data of 168 cultivated under near-zero growth conditions[50] are available from O. Kuipers.

### Sporulation and germination assays

To examine sporulation dynamics selected strains were cultivated in MSgg medium [51] at 30°C, 220 rpm, and total CFU and spore counts were analysed after 12, 24 and 36 hours. In sporulation assay performed with DTUB251 and DTUB254, the medium was supplemented with 0.2mM IPTG. To access the spore count, cells were incubated at 80°C for 20min, plated on LB-agar (1.5%) and the number of obtained colonies was divided by the number of colonies obtained prior to the heat-treatment. To access the germination, the culture incubation was prolonged to 72h to allow vast majority of cells to sporulate. Next, spores were washed 2× with 0.9% NaCl, and resuspendend in germination solution (0.6g KH2PO4, 1.4g K_2_HPO_4_, 0.2g (NH_4_)_2_SO_4_, 0.1g Na-citrate, 0.02g MgSO_4_×7H_2_O, 0.5g glucose, 3.56g L-alanine resuspended in 100ml of dH_2_O) to reach final OD_600_ cca 10. Decline of OD_600_ was measured immediately, indicating germination [52]. Additional assessment of germination dynamics was performed using real-time brightfield microscopy by inducing spores with L-alanine on agarose pads, as described previously [53]. Agarose pads (1.5%, 9 mm diameter, 1 mm height) were inoculated with 2.6 μl spore solution (3.75*10^5^ spores μl^−1^) and placed upside down into a 24-well glassbottom microtiter plate. Germination was induced by adding 5 μl of a 200 mM L-alanine solution to the top of the pad. Germination events were monitored by changes in grey level spore intensity. The fraction of germinated spores at time t was calculated as the number of germinated spores divided by the number of dormant spores before induction (i.e.by excluding pre-germinated spores).

### Spore selection experiment with NCBI 3610

Strains were cultivated in 10ml of MSgg medium in 100ml-glass bottles in 30°C with shaking at 220 rpm. Every 48 hours, three alternative transfer methods were applied: direct transfer of untreated cells to fresh medium, transfer of heat-treated cells (80°C for 20 min) and transfer of chemically treated cells (5% NaOH for 2 min, followed by washing in PBS). In each case, fresh cultures were initiated with 1% inoculum. Culture supernatants and cell pellets were collected prior each transfer to monitor phage release and genetic rearrangements, respectively. At each transfer, frozen stocks were preserved, to allow the analysis of subsequent steps of phage recombination in the future.

### Isolation of phage particles and phage DNA

Lysogens were cultivated in LB medium at 37°C with shaking at 200 rpm for 8h. Culture supernatants were collected, adjusted to pH of 7.0, filter-sterilized and mixed at a 1:4 rate with PEG-8000 solution (PEG-8000 20%, 116 g/l NaCl). After overnight incubation at 4°C, the solutions were centrifuged for 60 min at 12000 rpm to obtain phage precipitates. The pellets were resuspended in 1% of the initial volume in SM buffer (5.8 g/l NaCl, 0.96 g/l MgSO4, 6 g/l Tris-HCl, pH 7.5). Such phage solutions were visualized by transmission electron microscopy and used as a source of different phage variants, purified from single plaques. In plaque assay and further phage propagation from single plaques, Δ6 strain[54] was used as a host. Specifically, phage solutions were diluted in order to obtain well-separated single plaques. Selected plaques (differing with morphology) were carefully removed from the soft agar using sterile scalpel, resuspended in 200μl of SM buffer and used to infect exponentially growing phage-free host to allow propagation of selected phage variants. Phages were subsequently propagated in soft agar and liquid host suspension until the titer reached at least 10^9^ pfu/ml and then subjected to DNA isolation. Phage DNA was extracted using phenol-chloroform method, as described previously [55].

### Transmission electron microscopy

Before use, 400 mesh nickel grids with a 3-4 nm thick carbon film, CF400-Ni-UL EMS Diasum, were hydrophilized by 30 sec of electric glow discharging. Next, 5μl of purified phage solutions were applied onto the grids and allowed to adsorb for 1 minute. The grids were rinsed 3 times on droplets of milliQ water and subjected to staining with 2% uranyl acetate. Specifically, with a help of EM grid-grade tweezers, the grids were placed sequentially on droplets of 2% uranyl acetate solution for 10 sec, 2 sec and 20 sec. Excess uranyl acetate was wicked away using filter paper and the grids were allowed to dry overnight and stored in a desiccator until analysis. Transmission electron microscopy was performed utilizing a FEI Tecnai T12 Biotwin TEM operating at 120 kV located at the Center for Electron Nanoscopy at the Technical University of Denmark, and images were acquired using a Bottom mounted CCD, Gatan Orius SC1000WC.

### Prophage database construction and phage comparisons

*Bacillus* prophage database was constructed by finding genomic coordinates using Phaster software [56, 57] from fully assembled *Bacillus* genomes available at NCBI. All prophages between 80kB-50 kB, integrated between 1.9-2.3 Mb in the chromosome were selected for further analysis. Additional prophages, categorized as SPβ-like, were retrieved the genomes that gave BLAST hits to phi3T and SPβ, if these hits belonged to a prophage region that was at least 40kB. Duplicates and misassembled genomes (NZ_CP032855.1) were removed.

Interruption of *spsM* and *kamA* in all the selected lysogens was examined by genome BLAST against the sequence of an intact copy of these gene. All strains that carried a split copy of *spsM* and *kamA*, also carried a large prophage between left and right arms of these genes. In such cases, the Phaster-predicted terminal positions of the prophage was corrected to match the sequence splitting *spsM* or *kamA*. Integration genes of remaining large prophages were determined by extracting and clustering 1000bp-long prophage flanking regions using vsearch at 46% identity. These regions were then compared to well-annotated *B. subtilis* 168, using blastx, to find functional homologs.

The alignment of prophage sequences was performed in MAFFT program, using FFT-NS-2 algorithm and auto settings [58], phylogenetic tree was build using FastTree, using Jukes-Cantor model and auto settings [59, 60] and visualized in CLC Main Workbench. Phylogenetic tree of *B. subtilis* host strains was constructed using autoMLST (https://automlst.ziemertlab.com/) [61] based on 100 shared proteins. Two strains that were not lysogenic for SPβ-like prophage (MB9_B4 and MB9_B6) were included in the analysis to exclude SPβ prophage from the shared pool of proteins in the tree building. The three was visualized in CLC Main Workbench. Prophage annotation was performed using RAST annotation platform.

### Statistical analysis

Statistical differences between two experimental groups were identified using two-tailed Student’s *t*-tests assuming equal variance. Normality of data distribution was tested by Shapiro-Wilk and Kolmogorov-Smirnov tests. No statistical methods were used to predetermine sample size and the experiments were not randomized.

## RESULTS

### Strains evolved under a sporulation selection regime carry hybrid prophages

We previously showed that several passages of *B. subtilis* through the dormant spore stage leads to spontaneous release of phage particles into the culture medium (Fig. 1A). Spontaneous phage release occurred in *B. subtilis* 168 grown in rich medium under static, biofilm-promoting conditions after roughly 10 passages which involved selection for spores, but not in the absence of such selection in parallel populations [43]. Recently, similar spontaneous phage release was observed for *B. subtilis*NCBI 3610 grown in minimal medium under shaken-culture conditions after 10 passages through dormant spore stage [62]. Some of these phages resemble indigenous SPβ [63], but they facilitate killing of the ancestor strain [43] (Fig. 1B). To further characterise the phage-releasing evolved strains, three isolates [43] were subjected to genome sequencing (see Materials and Methods). *De novo* sequencing revealed the presence of an exogenous phi3T-like prophage (KY030782.1; 99.98% sequence identity) [64], which is closely related to SPβ (58% sequence identity) (Fig. 1C, Suppl. Fig. 1A, Suppl. Fig. 2A). We named this phage phi3Ts. The only difference between phi3T and phi3Ts was a 725 bp fragment (labelled ‘*s’* for sporulation-derived) within phi3Ts, replacing the 1265 bp fragment of phi3T (nucleotides 101,429-102,694; Fig. 1D). Although, the ‘s’ fragment shared no homology with phi3T or *B. subtilis* 168 chromosome, it could be found within SPβ-like prophages of six *B. subtilis* strains from different regions around the world (Fig. 1D). In the evolved strains, the phi3Ts prophage either disrupted the *kamA* gene located ~11 kb from SPβ, or it created a hybrid with SPβ with a ~11 kb fragment deleted between *kamA* and SPβ (Fig. 1C, Suppl. Fig 1A). In addition, sequencing coverage within the described prophage regions was increased severalfold, suggesting augmented replication of hybrid phage DNA (Suppl. Fig. 1B, Suppl. dataset 1).

**Figure 1.**
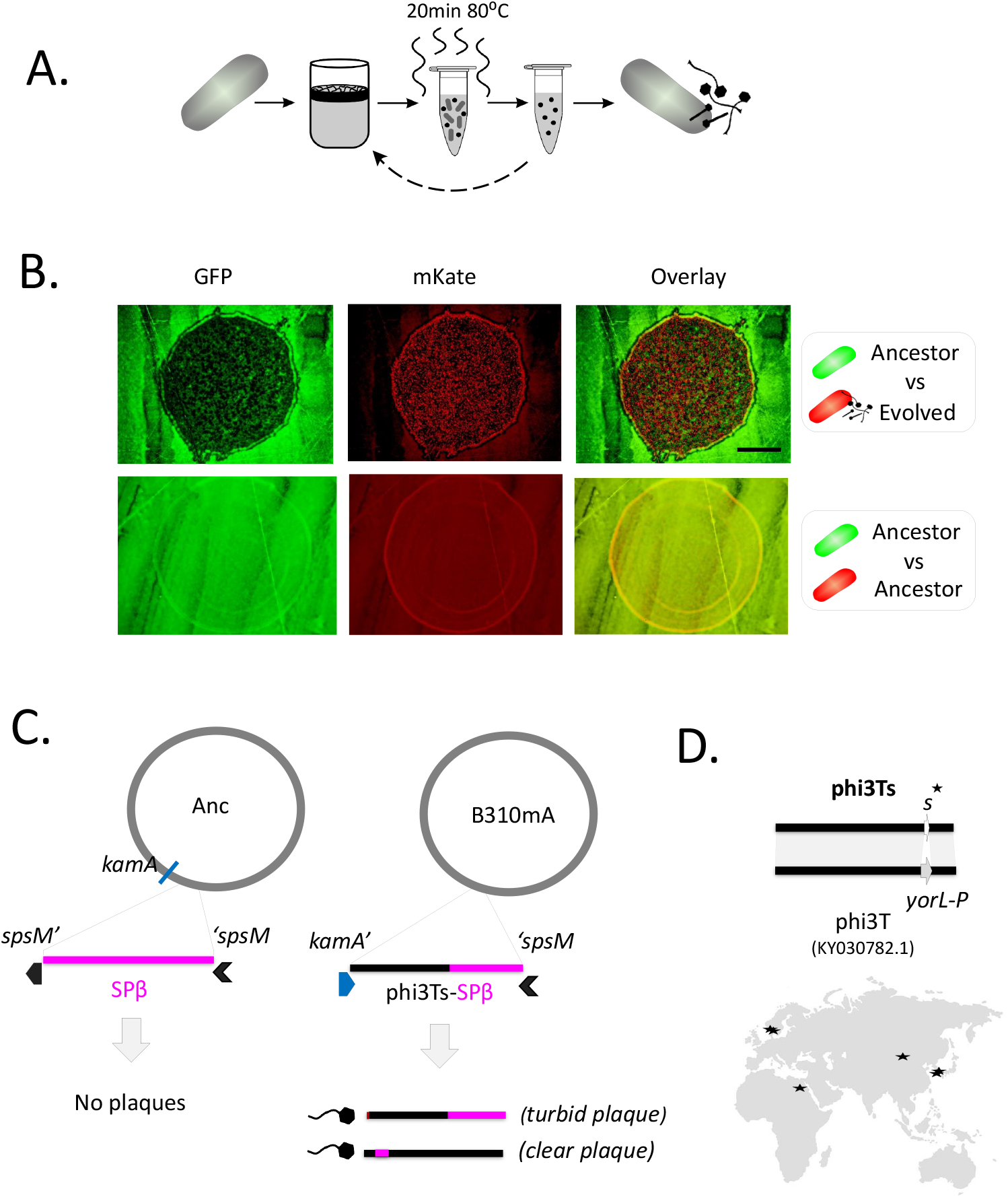
Changes within *B. subtilis* prophage sequence and integration site observed after prolonged sporulation selection regime. A) Experimental evolution with sporulation selection regime leads to spontaneous release of phage particles by the evolved strains[43]. B). Overnight culture of evolved *B. subtilis* strain B410mB *(amyE::mKate,* shown in red) was diluted 100’ and spotted on the lawn of undiluted *B. subtilis* ancestor strain *(amyE::gfp,* shown in green), resulting in a clearance zone, and growth of B410mB in that zone. The same experiment was performed using 100x diluted culture of ancestor strain *(amyE::mKate)* on a lawn of undiluted ancestor *(amyE::gfp),* as control.Scale bar=2.5mm. C) Schematic representation of genome rearrangements in one of the phage-releasing evolved strains (B310mA), compared to the ancestor (Anc). The evolved strains carry a hybrid prophage phi3Ts-SPβ. Fragments of phi3Ts are shown in black, while fragments of SPβ are shown in pink. Below, schematic representation of phage genomes, spontaneously released by B310mA. D) Schematic comparison of phi3Ts genome, with genome of Bacillus phage phi3T (KY030782.1). Fragment ‘s’ which is unique for phi3Ts, can be detected within prophage genomes of 6 *B. subtilis* strains, isolated in different parts of the world, specifically: SRCM103612 (South Korea), MB9_B1 and MB8_B1 (Denmark), JAAA (China), HMNig-2 (Egypt) and SSJ-1 (South Korea).

Comparison between phi3Ts and SPβ revealed that the first lacks putative repressor gene *yonR* as well as immunity-encoding gene *yomJ* what could explain spontaneous release of phage particles (Suppl. Fig. 2B).

### Strains lysogenic for hybrid prophages produce virulent hybrid phages

To examine which phage variants were released into the medium [43], phage-containing supernatants were spotted onto the lawn on Δ6 strain (derivative of 168 strain deprived of prophage elements [54]), followed by purification of phages from single plaques and sequencing (see Materials and Methods). Notably, each evolved strain produced a mix of turbid and clear plaques (Suppl. Fig. 1C). Turbid plaques are typical for temperate phages (like SPβ or phi3T), while clear plaques are usually formed by phages that have lost their lysogenic cycle [65]. Sequencing revealed that the spontaneously produced phages were either phi3Ts or phi3Ts-SPβ hybrids (Fig. 1C, Suppl. Fig. 1A, Fig. 2A). All phages obtained from the turbid plaques could also be identified within genomes of the evolved strains (Fig. 1C, Suppl. Fig. 1A, Fig. 2A). In addition, the genome of Hyb1^phi3Ts-SPβ^ (released by B310mA) was extended by a ~1.2 kb fragment of the host chromosome *(yozE,yokU,* and part of the *kamA* gene), indicating specialised transduction, a process that occurs when a phage picks up a fragment of host chromosomal DNA in the immediate vicinity of its attachment site (Fig. 1C, Fig. 2A). In contrast to the turbid plaque-creating phages, all phages obtained from clear plaques were phi3Ts-SPβ hybrids, which were not present on the chromosomes of their corresponding producers (Fig. 1C, Suppl. Fig. 1A, Fig. 2A) suggesting they were virulent phi3Ts-SPβ variants. We could also identify a variety of extrachromosomal phage DNA (epDNA) fragments ranging from 10.9 to 66 kb (Fig. 2B, Suppl. dataset 1, Suppl. Table 1). The epDNA was dominated by phi3Ts-SPβ recombinants, in which DNA from the two parental phages was joined at the homologous region (Fig. 2B). We identified 16 unique recombination sites (epDNA and hybrid phages combined), 15 of each were at regions of phi3Ts-SPβ homology, mostly focused around the left arm of phi3Ts/initial part of SPβ in *B. subtilis* chromosome (Fig. 2B., Suppl. Table 1). A single nonhomologous recombination site was found between a region following *yosA* gene of SPβ and unknown phi3Ts gene (Fig. 2B., Suppl. Table 1).

**Figure 2.**
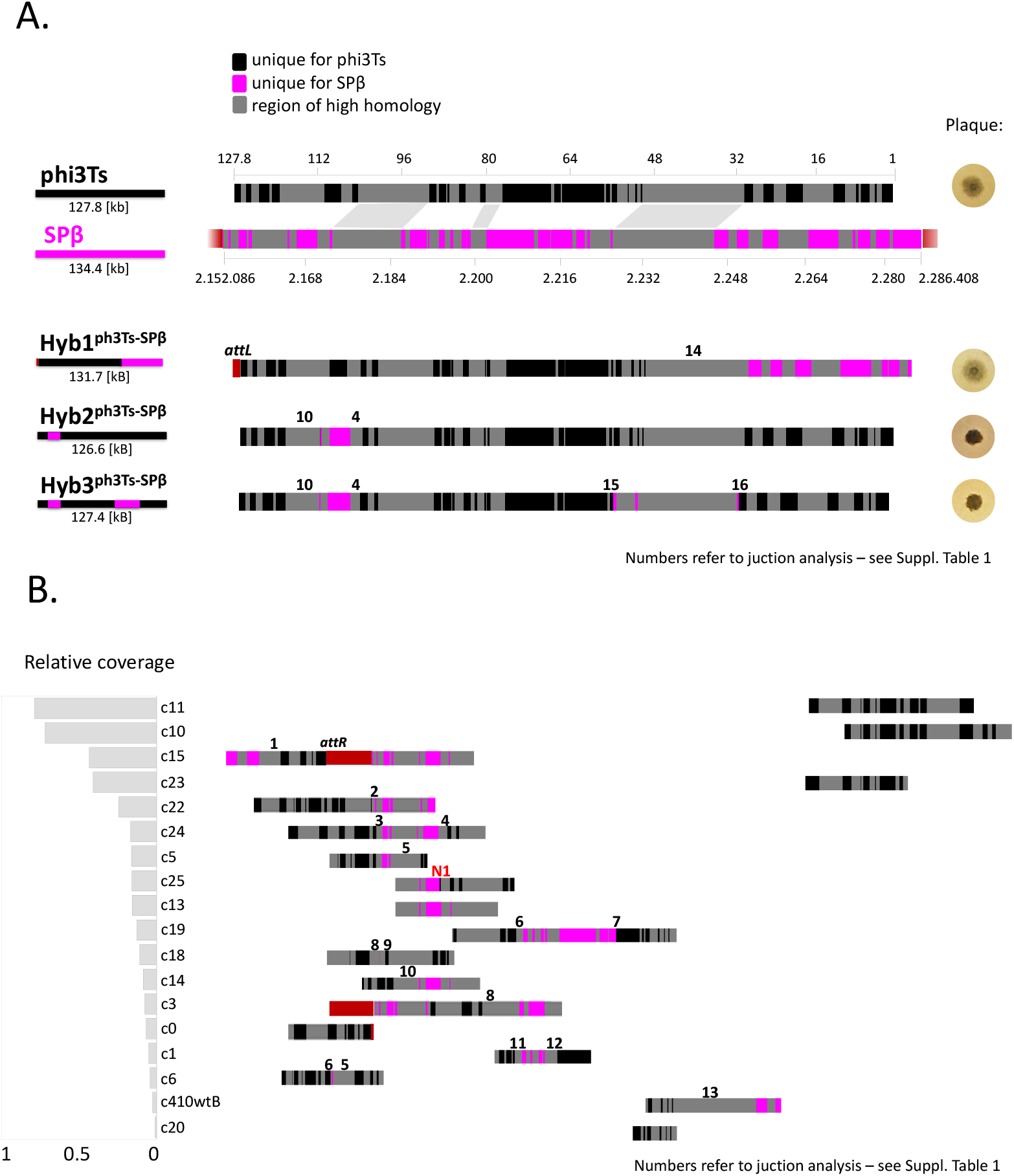
Hybrid phages and extrachromosomal fragments of phage DNA, detected in the evolved strains. A) Top: Genome comparison of phi3Ts and SPβ (Query cover=58%, Percent Identity=99.73%), where regions of high homology (73.6-100%) are shown in grey, and regions of 99% homology are connected. Segments that are unique for phi3Ts, or SPβ are highlighted in black and pink, respectively. Phage genomes are arranged according to their integration into the host chromosome, which is represented in red. hybrid phi3Ts-SPβ phages spontaneously released by the evolved strains. Below: Each hybrid phage contains DNA segments that are unique to SPβ or phi3Ts. Based on the position of these segments, the simplified schemes of recombinant phages were created (below the hybrid name). Hyb^phi3Ts-SPβ^ additionally carries fragment of the host chromosome, adjacent to the left att site. Picture of corresponding plaque is placed on the right site of phage maps. Numbers on phage maps indicate unique recombination sites, which are characterized in Suppl. Table 1. B) Extrachromosomal phage DNA fragments detected during PacBio sequencing, colored according to their homology to phi3Ts, SPβ, or fragments of host chromosome flanking phage integration sites. Fragments are ordered according to sequencing coverage relative to the chromosomal region, which is represented as bar chart on the left. Numbers on epDNA maps indicate unique recombination sites, which are characterized in Suppl. Table 1.

Although none of the hybrid epDNA was identical to sequences of hybrid phages released by the corresponding strains, certain homologous recombination sites in Hyb2^phi3Ts-SPβ^ and Hyb3^phi3Ts-SPβ^ were overlapping with these found in epDNA (Fig. 2, Suppl. Table 1). Finally, we also noticed that some epDNA fragments contained parts of the bacterial chromosome adjacent to the phi3Ts integration site, again pointing towards specialised transduction (Fig. 2B, Suppl. Table 1).

### A sporulation selection regime promotes foreign phage invasion

Next, we aimed to identify the source of phi3Ts DNA in the evolved host genomes. First, we repeated the mapping of raw sequencing reads from the *B. subtilis* 168 ancestral genome onto selected unique phi3T regions. Indeed, phi3Ts DNA was present in the ancestor strain at a very low but detectable level (Fig. 3, Suppl. Fig. 3A). On the other hand, phi3T could be clearly detected by mapping of sequencing reads of the evolved strains (Suppl. Fig. 3A).

**Figure 3.**
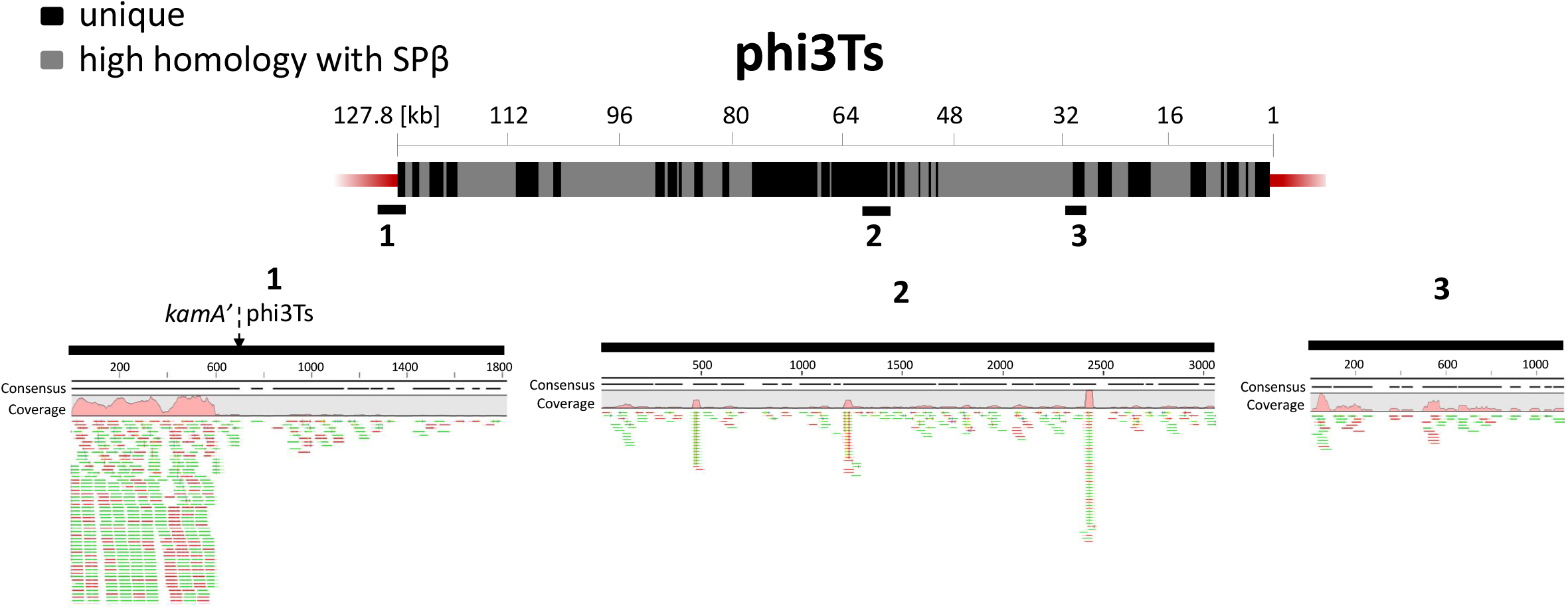
Detection of phi3Ts DNA in the ancestor strain *B. subtilis* 168 through mapping of raw sequencing reads. Top: Representation of phi3Ts genome according to its homology to SPβ prophage. Fragments of high homology to SPβ (73.6-100%) are shown in grey, while fragments that are unique to phi3Ts are shown in black. Bars 1,2 and 3 correspond to DNA sequences that are unique for phi3Ts and that were used as targets for raw reads mapping (lower part). Green and red bars represent reads obtained from forward and reverse strands, respectively.

This result was additionally confirmed by PCR (Suppl. Fig. 4). As expected, the evolved strains B310mA and B410wtB, where integration of phi3Ts into *kamA* locus was confirmed by genome sequencing (Fig. 1C, Suppl. Fig. 1A), were negative for intact *kamA,* while B410mB gave a weak product, despite similar integration of phi3Ts (Suppl. Fig. 1A), what could be explained by incorporation of *kamA* into its epDNA (Fig. 2B, Suppl. Fig. 4A).

The above analysis indicates that low copy number phi3Ts was present in the *B. subtilis* 168 stock from the start. Since *B. subtilis* 168 has been shared among research labs around the world, phi3Ts could also be ‘hiding’ in culture stocks of other research labs. Accidental detection of such low copy number phage DNA is nearly impossible, because (i) sequencing reads matching phi3Ts would be filtered out during standard mapping pipelines, and (ii) phi3Ts appears to only multiply and manifest itself under specific selection regimes. To check for possible contamination of other *B. subtilis* stocks with phi3Ts, we mapped raw sequencing data available in the NCBI database to the phi3T genome (KY030782.1), but found no evidence of phi3Ts contamination (Suppl. Fig 3B). We also PCR-screened a larger collection of *B. subtilis* 168 stocks from different labs around the world[66] for the presence of phi3Ts and obtained a very strong positive result for *B. subtilis* 168 ‘Newcastle’, suggesting that this strain was lysogenized by phi3Ts, or a very similar prophage (Suppl. Fig. 4B). We also confirmed that, the Newcastle 168 strain contained the ‘s’ fragment, a unique sequence allowing the phi3Ts phage to be distinguished from the previously sequenced phi3T (Fig. 1D).

As phi3Ts multiplies under a prolonged sporulation selection regime, we contacted colleagues who also performed experimental evolution with *B. subtilis* strains imposing the same or similar selection [45, 50]. A group from the University of Wisconsin-Madison, agreed to share raw sequencing data obtained from 12 evolved single isolates. We did not find any mutations within prophage regions (Suppl. dataset 2) nor phi3T-specific DNA fragments in the raw sequencing data (see Materials and Methods; Suppl. Fig. 5).

We also approached a group from the University of Groningen, who performed experimental evolution of *B. subtilis* 168 under nutrient-limited conditions in which bacteria could neither grow nor complete sporulation (due to *sigF* deletion) [50]. Mapping their raw sequencing reads to the phi3T genome clearly revealed the presence of phi3T-specific reads (Suppl. Fig. 3C). Similar to our case (Fig. 3, Suppl. Fig. 3A), the phage DNA was already present at the start, and it either gradually decreased or increased in two different biological samples (Suppl. Fig. 3C).

The above results suggest that the prophage activation scenario requires not only a sporulation selection regime, but also contamination with low copy number phi3Ts DNA or phage particles. The exchange of strains between Newcastle University (the origin of *B. subtilis* 168 PCR-positive for the phi3Ts-specific fragment) and the University of Groningen, and later between the University of Groningen and our lab, represents a possible transmission route for phi3Ts.

Finally, the evolution experiment performed previously [43] was repeated under the sporulation selection regime using the undomesticated *B. subtilis* NCIB 3610 (hereafter 3610) strain in which the presence of phi3Ts DNA could not be detected during analysis of genome sequencing (Suppl. Fig. 3D, Suppl. Fig. 6A) or by PCR (Suppl. Fig. 6B). To our surprise, lytic activity (Suppl. Fig. 7A) and the release of phages (Suppl. Fig. 7) were observed as early as the fourth transfer when the sporulation selection regime was applied. Similarly, targeted PCR analysis of host DNA revealed a gradual increase in the phi3T-specific PCR product and a gradual decrease in the PCR product corresponding to intact *kamA* (Suppl. Fig. 8). This experiment again demonstrated unsupervised outspread of phi3Ts, taking place solely in populations subjected to sporulation/spore revival selection regime. The latter either facilitates contamination by phi3Ts or propagation of this phage from levels that are beyond PCR or sequencing detection limits.

### Foreign phages modulate sporulation dynamics

We next explored whether propagation of low copy number phi3Ts DNA and its integration into the *kamA* gene has any positive fitness effects on *B. subtilis.* Since expression of *kamA* is dramatically increased upon sporulation entry (http://subtiwiki.uni-goettingen.de), we hypothesised that *kamA* may encode a product that is metabolically costly and/or toxic for the bacterium, hence strains lysogenized by phi3Ts/phi3Ts-SPβ hybrid prophage lysogen may benefit from inactivation of this gene (Suppl. Figure 9A). However, competition assays between wild-type vs. *ΔkamA* strains with and without sporulation selection revealed no difference in performance between strains (Suppl. Fig. 9B).

Next, we examined the sporulation and spore revival dynamics of *B. subtilis* 3610 lyzogenized by phi3T, available from the Bacillus genetic stock center (BGSC). We observed that the phi3T lysogen sporulated prematurely compared with the wild-type strain (Fig. 4). We also observed a general trend indicative of better revival of the phi3T lysogen (Suppl. Fig. 10A), which may include contributions from faster germination (Suppl. Fig. 10B) and/or an altered frequency of premature germination during dormancy (Suppl. Fig. 10C). These results suggest that the spread of phi3Ts might be partly driven by host benefits from the regulatory arsenal associated with phi3Ts and its hybrids.

**Figure 4.**
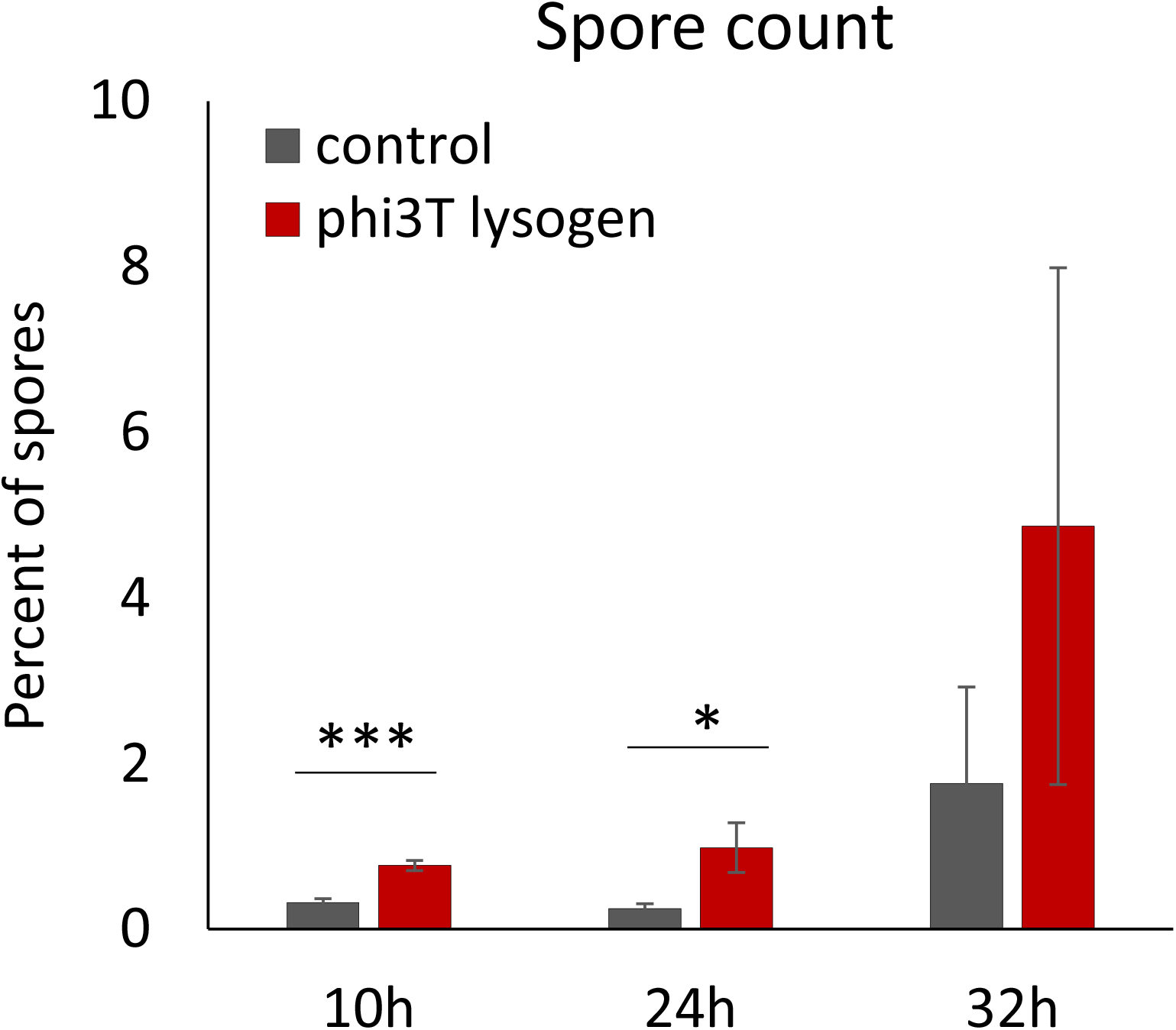
Effect of phi3T lysogenization on *B. subtilis* sporulation and germination dynamics. A) Sporulation dynamics. Percentage of spores compared to total cell count, were examined in *B. subtilis* 3610 and the same strain lysogenized by phi3T phage, in 3 different time points of growth in minimal medium (MSgg). Data represent an average from 4 biological replicates, error bars correspond to standard error.

Notably, sporulation regulators have been previously linked to mobile genetic elements (MGEs) in this species [67–69]. Annotation of phi3Ts and phi3Ts-SPβ hybrids (see Materials and Methods) revealed the presence of several genes that could modulate sporulation or spore traits. Specifically, we found a gene (labelled as *rapX)* encoding a putative Rap phosphatase (Suppl. Fig. 2A, Suppl. Table 2) sharing high amino acid sequence identify with RapA (unique for phi3Ts) that is known to modulate sporulation timing [70]. We also found that the ‘s’ phi3Ts marker sequence may encode stationary phase survival protein YuiC (100% confidence Phyre prediction = 100% confidence), hence we labelled this sequence *spsX* (Suppl. Fig. 2A). Indeed, overexpression of *spsX* from ectopic locus resulted in small but significant increase in percent of early spores as well as faster germination of *B. subtilis* NCBI 3610 (Suppl Fig. 11).

In addition, we identified *sspC* that controls spore resistance traits and encodes an acid-soluble protein involved in spore DNA protection (present on both SPβ and phi3Ts) [71]. Therefore, phi3T/phi3Ts may encode proteins that influence the *B. subtilis* life cycle during sporulation and spore revival.

### Recombination between SPβ-like prophages takes place on global and local ecological scales

To understand the ecological relevance of extensive phage recombination observed under a sporulation selection regime, we analysed prophage elements within the *B. subtilis* clade including *B. cereus* for comparison of more distant species (see Materials and Methods) (Suppl. dataset 3). We immediately identified a cluster of rather large (>100 kb) prophages integrated close to the replication terminus, just like phi3Ts, SPβ or phi3Ts-SPβ hybrids. These large prophages were found in 78 out of a total of 320 genomes, mainly (86%; 67/78) within representatives of *B. subtilis* (34/78), *B. velezensis* (17/78). *B. amyloliquefaciens* (10/78) and *B. licheniformis* (6/78) species (Fig. 5A, Suppl. dataset 3, Suppl. Fig. 12).

**Figure 5.**
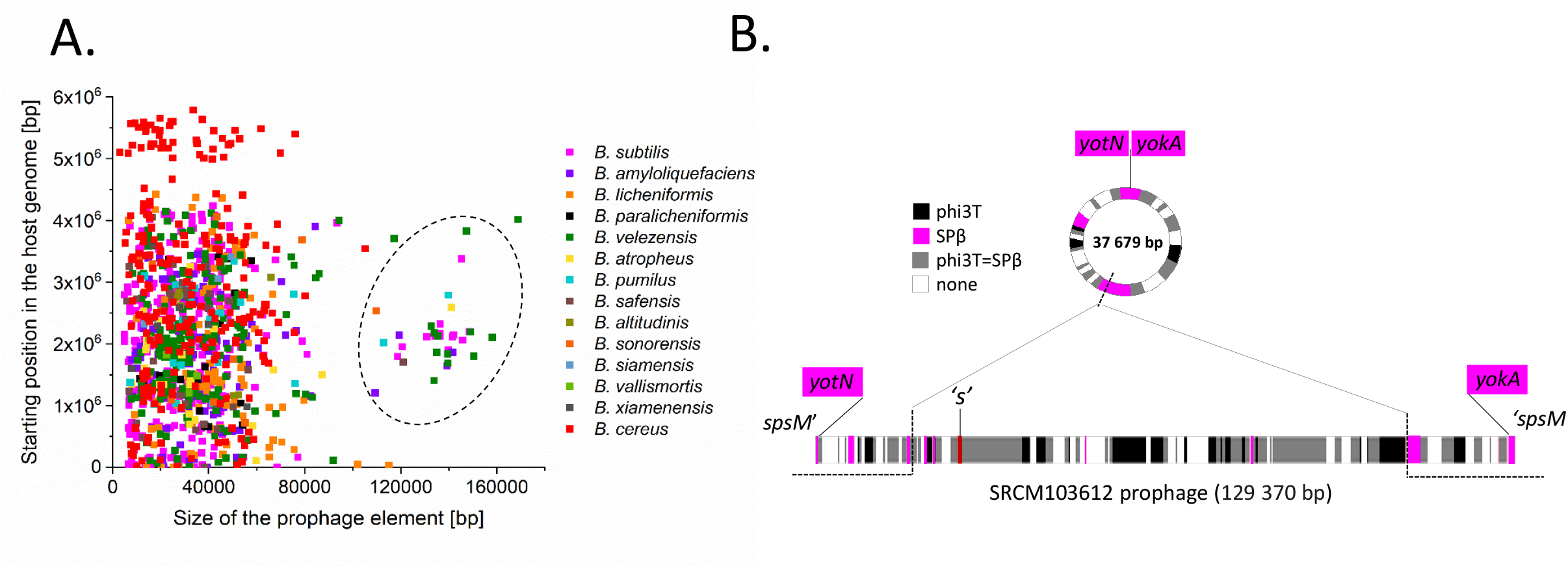
Overview of prophage elements of natural *Bacillus* sp. isolates. A) Prophage elements were extracted from fully assembled genomes of *Bacillus* sp. and plotted according to size and integration position in the chromosome. Cluster of large prophages, integrated in the area of replication terminus could be detected (black dotted line). B) Schematic representation of SPβ-like prophage found in *B. subtilis* SRCM 103612, isolated from traditional Korean food. The prophage genome was colored according to its homology to phi3Ts and SPβ. Extrachromosomal phage DNA found in this strain is matching left and right arms of the chromosomal prophage.

We further identified 23 strains in which large prophages split the *spsM* gene in a manner identical to SPβ (Suppl. dataset 3), and four *B. subtilis* isolates in which the *kamA* gene was split by a prophage at exactly like observed in the our evolved strains lysogenized by hybrid phages (Suppl. dataset 3). In the remaining Bacillus strains, the large prophages were mostly integrated close to sporulation-related genes (Suppl. dataset 3). Interestingly, 10 strains carried extrachromosomal phage DNA (as predicted by Phaster; Suppl. dataset 3), and in one of them (*B. subtilis* SRCM103612) this epDNA was a truncated version of an SPβ-like prophage present within the chromosome (Fig. 5B). The SRCM103612 prophage contained regions sharing homology to both SPβ and phi3Ts, indicating recombination and an unstable lysogenic cycle within SPβ-like recombinant phages in natural *B. subtilis* isolates (Fig. 5B).

To access the natural diversity of large SPβ-like prophages, we collected *Bacillus* sp. genomes carrying a large prophage splitting *spsM* or *kamA*. Notably, such prophage was present in roughly 39% of all unique *B. subtilis* genomes available (28 out of 72). We compared the phylogenetic tree obtained for these host strains (see Materials and Methods) with the phylogenetic tree obtained for their SPβ-like prophages (Fig. 6AB). The strains could be divided into six phylogenetic clades (Fig. 6A), while prophages clustered into three clades: ‘conservative’, ‘hybrid’ and ‘diverse’. The ‘conservative’ clade comprised prophages that were nearly identical to SPβ, all residing within closely related *B. subtilis* strains (Fig. 6AB). The ‘hybrid’ clade comprised phi3T, phi3Ts and all phi3Ts-SPβ hybrids that evolved in the above described experiments (Fig. 6B). Within the ‘diverse’ clade the prophage relatedness did not match the phylogenetic relatedness of the hosts (Fig. 6AB). Here, we found that among *B. subtilis* isolates from the same soil sample below the mushroom [72], one strain (MB8_B7) carried an *spsM*-integrated SPβ prophage, one strain (MB8_B1) carried a SPβ-like prophage in *spsM,* and one strain (MB8_B10) carried an SPβ-like prophage in *kamA* (Fig. 6AB). Notably, all members of the ‘conservative’ clade carried an intact copy of ICEBs1 that was shown to block the SPβ lytic cycle [44], while this element is missing in all members of the ‘diverse’ clade (Fig. 6B). Like phi3Ts and phi3Ts-SPβ hybrid phages, several representatives of the ‘diverse’ cluster lacked the putative repressor gene *yonR,* which was present in all members of the ‘conservative’ cluster (Fig. 6B, Suppl. dataset 3). Finally, we could clearly see modules sharing high homology with SPβ and phi3T in the large prophages (Fig. 6C). These results are consistent with our lab data showing that SPβ-like phages diversify in nature, and this diversification may be constrained by other MGEs present on the host chromosome and facilitated by loss of repressor gene through homologous recombination.

**Figure 6.**
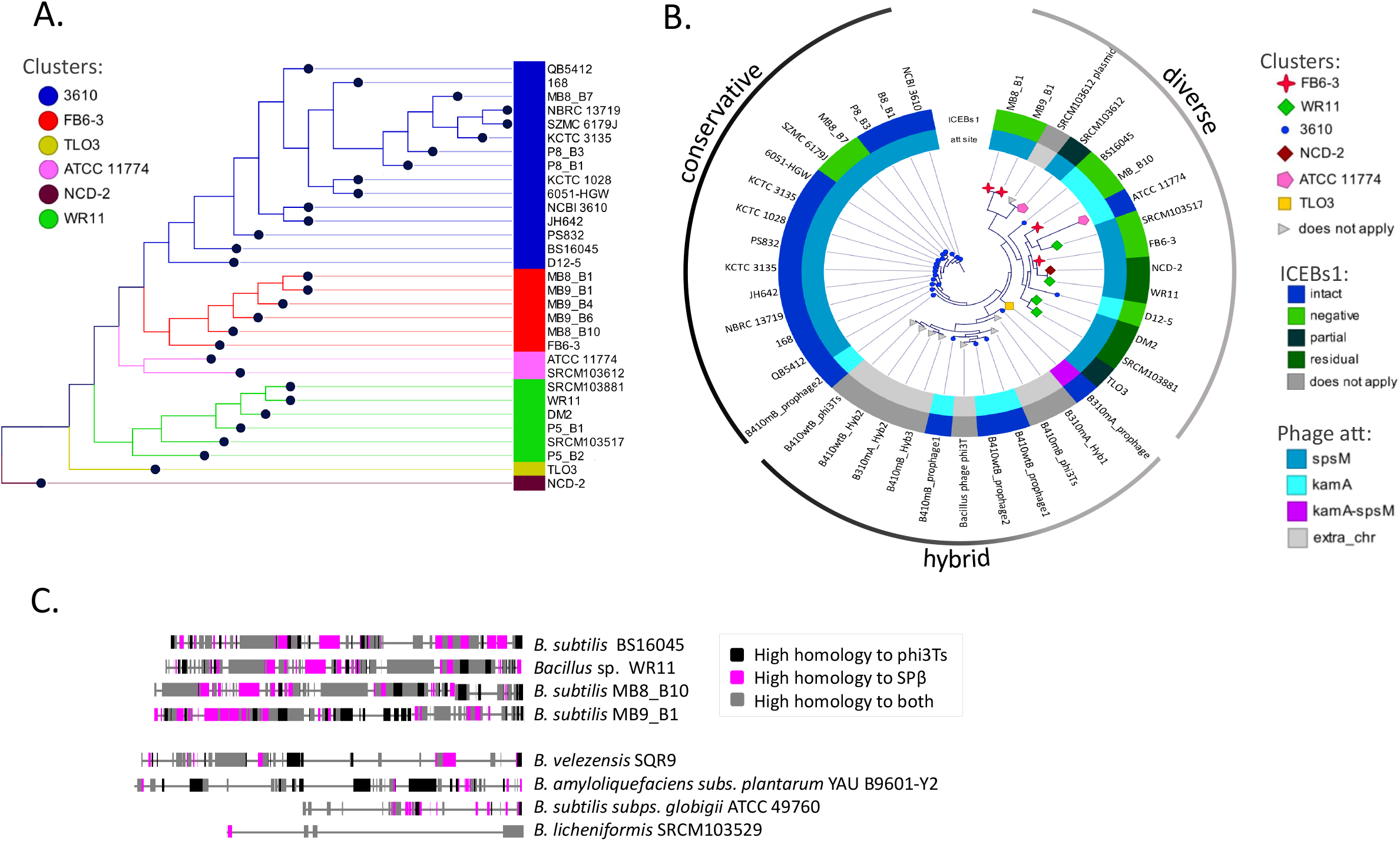
Natural diversity of SPβ-like phages. A). Phylogenetic tree of *B. subtilis* strains that carry SPβ-like prophage in *spsM* or *kamA* gene, and two control strains that are free from such prophage. The tree was arbitrarily divided into 6 clades. B) Phylogenetic tree of SPβ-like prophages hosted by the strains in A). Inner circle shows prophage integration site, while outer circle indicates presence/absence of conjugative element ICEBs1, which blocks SPβ lytic cycle C). Selected prophages of Bacillus sp. colored according to their homology to phi3T and SPβ. The upper 4 sequences integrate either in *kamA* or *spsM* and clearly belong to SPβ-like phages. Bottom four sequences come from other *Bacillus* species, and although they are more distant to phi3T or SPβ, they still carry segments of high homology with these phages. Explanation of ICEBs1 figure legend: intact – intact copy (100% identity to *B. subtilis* 168 or NCBI 3610) of ICEBs1 conjugative element is present; negative – lack of BLAST hits to ICEBs1 sequence; partial – at least 70% of ICEBs1 sequence is present; residual – less than 5% of ICEBs1 sequence is present.

## Discussion

The importance of phages in the ecology and evolution of bacteria is indisputable. Genome comparison suggests that prophage elements undergo pervasive domestication within their hosts that gradually lose the ability to reproduce via the lytic cycle [73]. Our current work demonstrates an opposite scenario, where after a prolonged sporulation/spore revival selection regime, a latent prophage of *B. subtilis* (SPβ) [43, 44] regains its lytic reproductive cycle via recombination with ‘foreign’ phage DNA (phi3Ts). Prophage recombination did not occur in the absence of sporulation/spore revival selection regime, as evident from our previous experimental evolution studies [74, 75], as well as this work (Suppl. Fig. 7).

The fact that phi3Ts only manifests itself under specific conditions (a prolonged sporulation/spore revival selection regime) is reminiscent of previously described examples of *Proteobacteria* phages [76, 77]. Specifically, the lytic phage SW1 can thrive undetected within *E. coli* populations, but manifests itself in spontaneous plaque formation after overexpression of a putative methylase from an indigenous cryptic prophage [76]. Likewise, lytic variants of P22 spontaneously form upon purine starvation of the *Salmonella typhimurium* host [77]. It was later realized that P22 coexists with its host in so called carrier state, when extrachromosomal phage is asymmetrically inherited by only one of the sibling cells after cell division, resulting in coexistence of resistant and susceptible host populations [78, 79]. Similar mechanism could maintain low amount of phi3Ts within *B. subtilis* population.

Identification of phi3Ts and phi3Ts-SPβ hybrids within *B. subtilis* populations evolved under sporulation/spore-revival selection regime, sheds new light on our previous work, that changes within SPβ regions specifically promote the spread of biofilm-deficient strains in the population [43]. Present study reveals that while biofilm-proficient strain (B410wtB) still carries intact phi3Ts and SPβ at different loci, biofilm-deficient strains (B310mA and and B410mB) are lysogenized by phi3Ts-SPβ hybrids, what also results in genome deletion (Fig. 1C, Suppl. Fig. 1A), offering a potential explanation for fitness advantage of these strains. It remains to be tested whether biofilm-deficiency is linked to the likelihood of phi3Ts-SPβ recombination.

Exactly how sporulation promotes the spread of ‘foreign’ phage and its hybrid derivatives requires further molecular studies. There are two, not mutually exclusive, hypotheses: (a) induction of the phage lytic cycle in a small fraction of sporulating cells leads to rapid spread of phi3Ts within the population, infection of phi3Ts-susceptible sporulating cells, segregation of phage DNA into forespores, and trapping of many of its copies in spores, followed by the release of phages upon germination, as observed previously for lytic *B. subtilis* phages [80, 81]; (b) since phi3T lysogeny (KY030782.1; 99.98% sequence identity with phi3Ts) [64] results in earlier sporulation and potentially improved spore quality, integration of this phage into the chromosome may be adaptive for the host. As the functions of most phi3Ts genes are obscure, it is difficult to identify the potential phage-encoded regulatory genes that could affect the host life cycle. We show that *spsX* gene could be one possible candidate modulating sporulation and spore revival timing. Another suspect was *rap-phr* cassette (matching *rap* present in phi3T and phi3Ts) that have been previously found within other MGEs of *B. subtilis,* and have been shown to modulate the timing of sporulation [67–69], however we could not detect significant effects of *rapX* overexpression on sporulation/spore revival timing, contrary to recently hypothesized function of this gene in phi3T [82]. Phi3Ts phage genes (e.g. *sspC)* could modulate the production of resistant and viable spores [71] and/or reduce sporulation failure and premature germination [83]. In addition, the spore revival traits of lysogens may also be indirectly affected by the modulation of sporulation timing [53]. Whatever the exact molecular mechanism and its evolutionary driver, the activation of latent prophage elements upon sporulation/spore revival treatment expands the intriguing connections between sporulation of *Firmicutes* and phages infecting and lysogenizing these species [31, 37, 42, 84–86].

Based on comparison of the sizes and integration sites of prophage elements within Bacillus sp., SPβ and phi3T clearly belong to a distinct prophage group. These phages appear extremely large (2-3-fold larger than average prophages), and they possess sophisticated communication systems that are potentially capable of sensing the frequency of lysogenized hosts [64, 87] or biosynthetic gene clusters [88, 89], and functional genes related to host dormancy [90, 91]. A high level of homology between SPβ and phi3Ts offers extensive regions for homologous recombination, which can be additionally promoted by the recombination machinery involved in natural competence [92] and in non-homologous end-joining repair [93]. It appears that the absence of other mobile genetic elements (e.g. ICEBs1) constraining the phage lytic cycle [44] may also correlate with a higher level of phage diversification. It is possible that such phage recombination could be facilitated by regulatory excision of SPβ from the chromosome in the sporulating mother cell [31, 94].

Our work shows how large SPβ-like prophages of *Bacilli* pervasively recombine during sporulation, providing new model system to study bacterial evolution in which phages serve as an evolutionary driving force. These observations shed new light on the interplay between bacteria and their phages; while temperate phages commonly undergo domestication [73], they may easily regain genetic mobility by recombination with other phages, thereby altering the physiology, social interactions and evolution of their hosts. The crucial impact of prophage elements on ecological interactions within closely related strains has already been demonstrated for other species [7, 76, 95, 96]. Herein, we showed that such antagonism emerges during the early steps of phage diversification, which suggesting that speciation of prophage elements may be the first step toward speciation of host bacteria.

## Acknowledgements

The authors thank M. Kilstrup, P. Sazinas, K. Middleboe, D. Castillio and P. Stefanic for their valuable comments. We are profoundly grateful to O. Kuipers, A. de Jong and W. Overkamp from University of Groningen, for sharing their raw sequencing data and all relevant information, which allowed us to finalize the manuscript. This project has received funding from the European Union’s Horizon 2020 research and innovation programme under the Marie Skłodowska-Curie grant agreement No 713683 (H.C. Ørsted COFUND to A.D.), Individual grant from Friedrich Schiller University Jena to support postdoc researchers to A.D., and supported by the Danish National Research Foundation (DNRF137) for the Center for Microbial Secondary Metabolites. Funded in part by NIH R01GM121865 to BMB.

## Authors contributions

AD and ATK designed the study. AD, PB, ZH, CK performed experiments. AD and MLS performed bioinformatics analysis. PK performed electron microscopy, GM performed genome sequencing and analysed the data, BB and BMB shared sequencing data. AD wrote the manuscript. All authors contributed to final version of the manuscript.

