## Supplementary figures for "Pervasive prophage recombination occurs during evolution of spore-forming *Bacilli*"

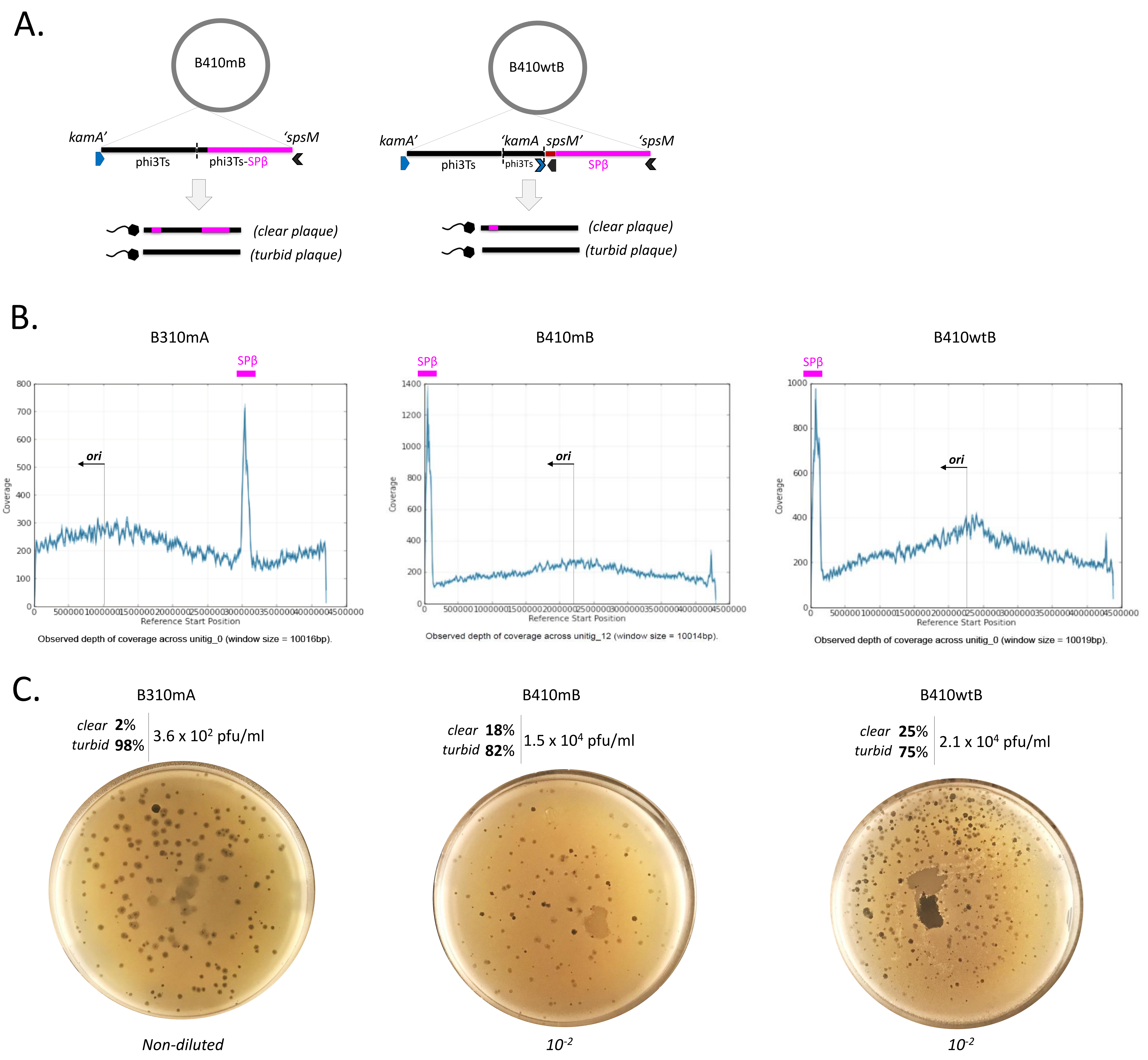


Supplementary Figure 1. Changes within *B. subtilis* prophage region upon sporulation selection regime and spontaneous phage release. A) Schematic representation of genome rearrangements the phage-releasing evolved strains (B410mB and B410wtB). The evolved strains carry a phi3Ts prophage and a phi3Ts-SPβ hybrid (in case of B410mB), or phi3Ts and its truncated variant integrated into *kamA* gene and an intact copy of SPβ (in case of B410wtB). Fragments of phi3Ts are shown in black, while fragments of SPβ are shown in pink. Below, schematic representations of phage genomes, spontaneously released by B410mB and B410wtB. B) Genomic coverage data obtained after sequencing and de novo assembly of evolved strains using PacBio method. Origin of replication was determined based on previously published genomes. Increased coverage within SPβ region was highlighted. C) Plaques obtained from PEG-8000 precipitated supernatants of B310mA, B410mB and B410wtB on lawn of prophage-free strain Δ6. Each precipitate produced different relative amounts of clear and turbid plaques.

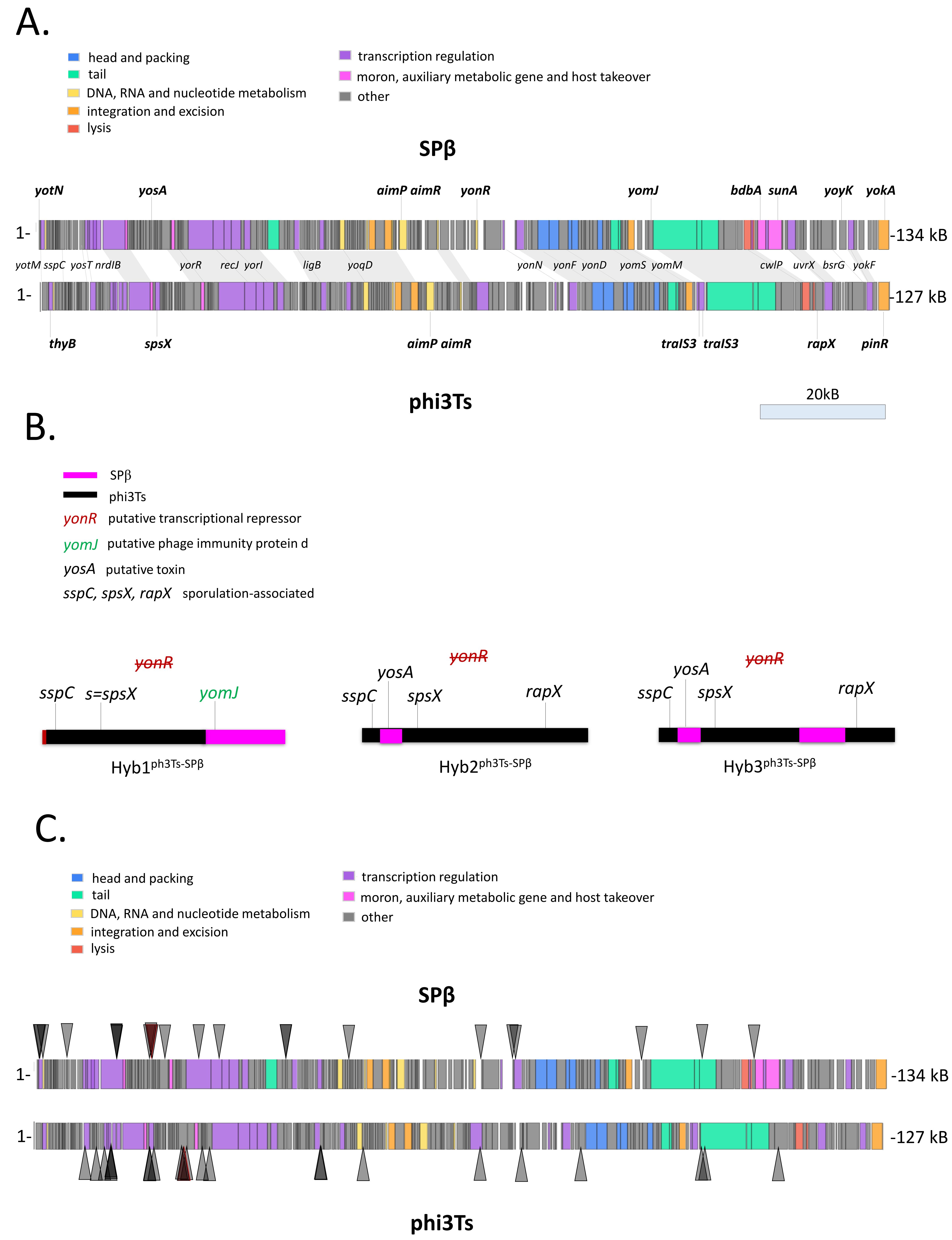


Supplementary Figure 2. Functional genes in SPβ, phi3Ts and hybrid phages. A) RAST annotation software was used to annotate phi3Ts in order to extract potential functional differences between phi3Ts and SPβ, as well as potential functional contribution of both phages to hybrid phage DNA. Phage maps with coloring acc. to functional category were retrieved from PhROGs website. Shared functional genes are shown in between the two maps, genes that are unique for SPβ are above SPβ map and genes that are unique for phi3Ts are shown below phi3Ts map. Putative functions of all genes are described in Supplementary Table S2. B) Schematic representation of hybrid phages, with potentially important genes coming from SPβ or phi3Ts. All hybrids contain *sspC* encoding for soluble acid protein, important for spore DNA protection, as well as *spsX* (referred to as sequences in Fig1), which may encode for stationary phase survival protein. In addition, neither phi3Ts nor the hybrids carry *yonR*, which is a putative phage repressor, which could explain unstable lysogeny. C) Maps with indicated recombination hotspots, which were determined based on Supplementary Table 1.

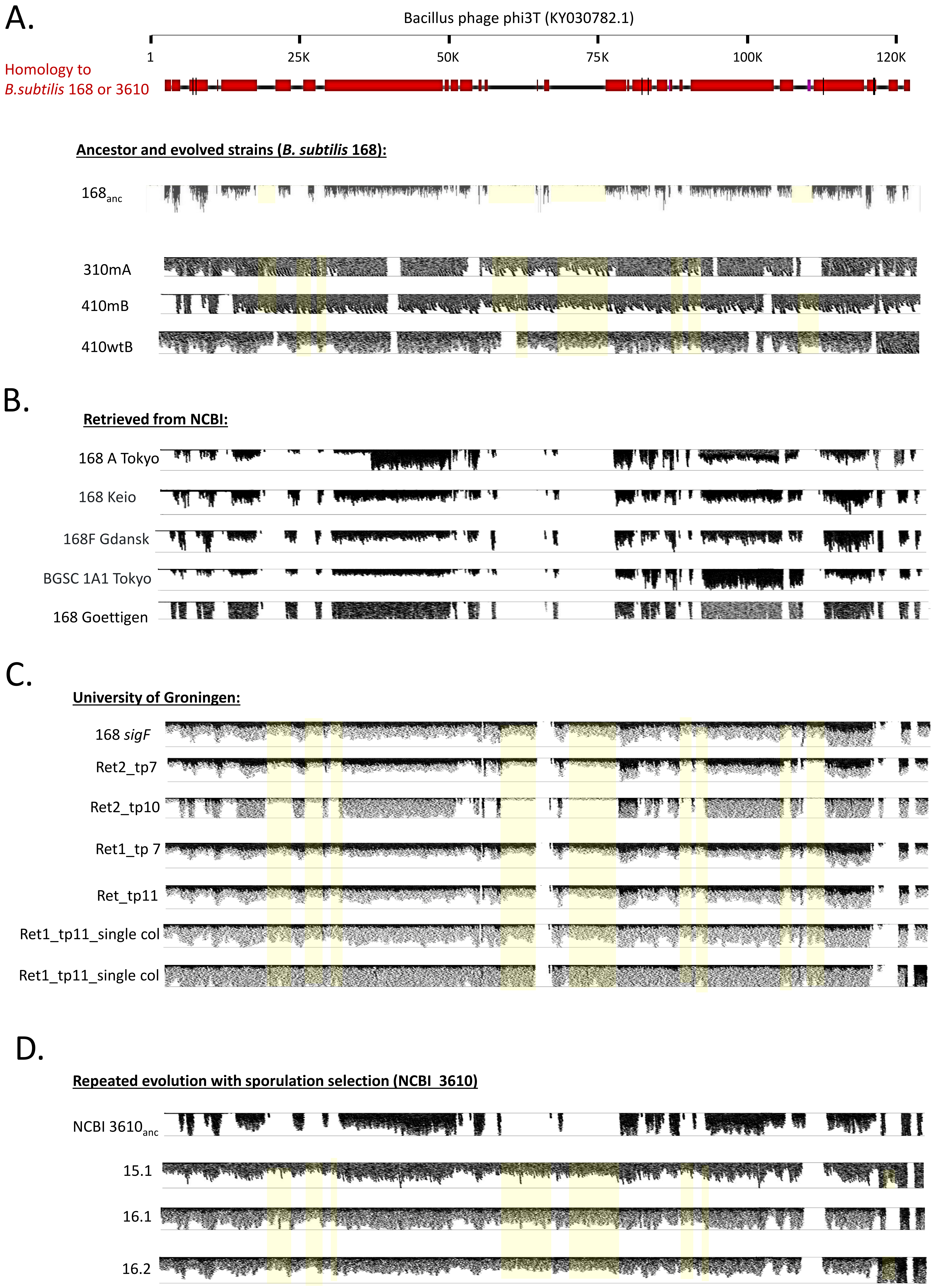


Supplementary Figure 3. Detection of phi3T DNA fragments in *B. subtilis* strains. Raw Illumina (or Solid for 168anc) reads from the sequencing several different isolates were mapped onto DNA of phi3T (KY030782.1) using Bowtie2 package in Galaxy platform (https://cpt.tamu.edu/galaxy-pub) and visualized using Trackster tool. Selected reads that overlap with unique phi3T fragments were highlighted in yellow. The strains that were examined include: A) *B. subtilis* 168anc used in this project and the evolved strains 310mA, 410mB and 410wtB; B) Sequenced genomes of 168 deposited at NCBI by different research groups, specifically, 168 A Tokyo (DRX007636), Keio (DRR000002), 168F Gdansk SRR1663459, BGSC 1A1 Tokyo DRR008456, 168 Goettigen (RR7967955); C) Sequencing reads of *B. subtilis* Δ*sigF* (ancestor) and strains evolved under near-zero growth conditions51, shared by colleagues from University of Groningen. Ret1 and Ret2 stand from Retentostat 1 and Retentostat 2 while tp correspond to different timepoints; D) Sequenced genome of NCBI 3610 and its 3 derivatives that evolved under sporulation selection regime.

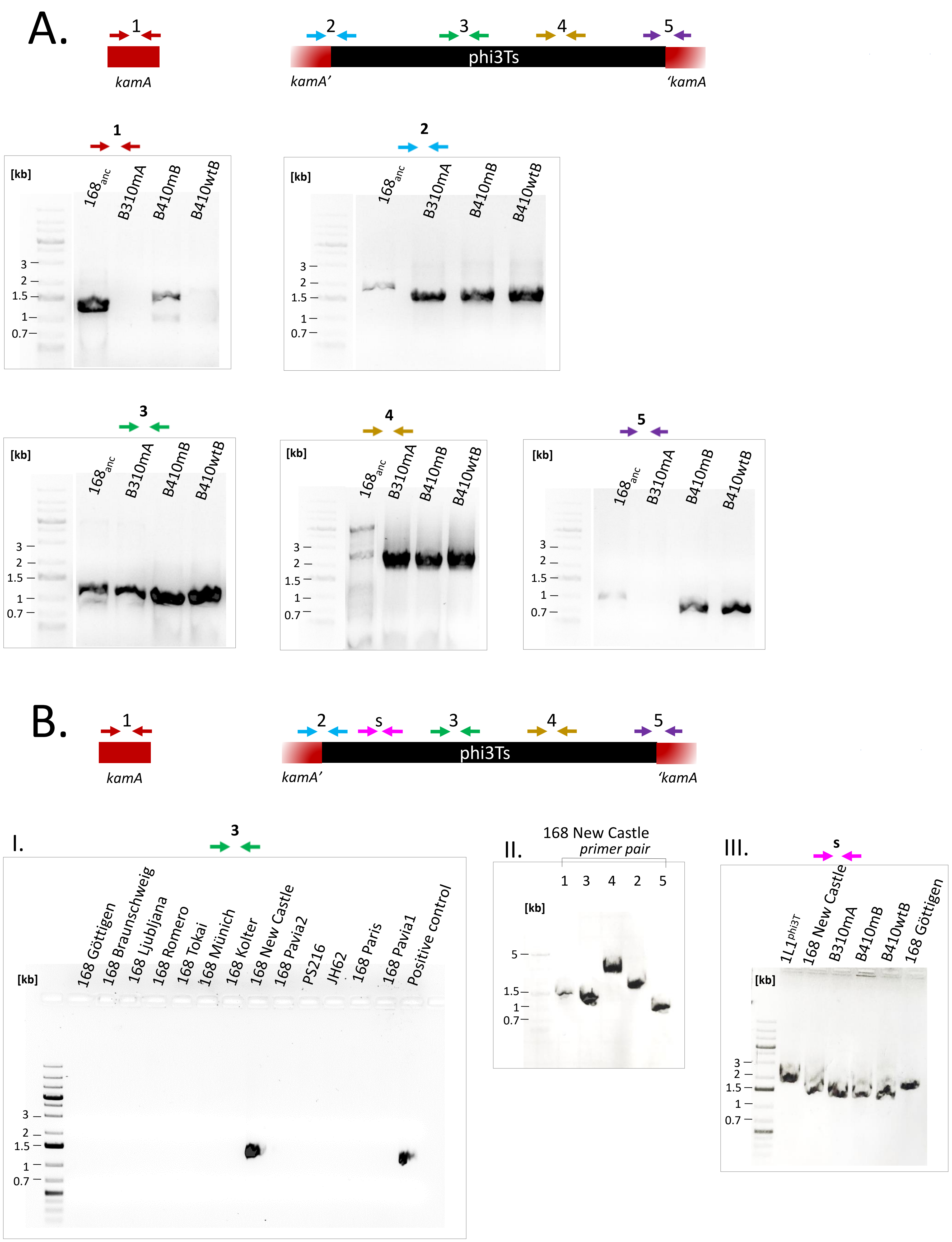


Supplementary Figure 4. Evaluation of the ancestor and phage-releasing *B. subtilis* by PCR. A) Ancestor strain and the three evolved strains were evaluated for presence of intact copy of *kamA* gene, phage integration into *kamA* and two unique sequences of phi3Ts, using five primer sets. Primer pairs used and expected molecular mass of the PCR product were as follows: 1 – oAD36/oAD37, 1366 bp; 2- oAD42/oAD43, 1819 bp; 3-oAD38/oAD39, 1121 bp; 4- oAD40/oAD41, 3033 bp; 5- oAD44/oAD45, 901 bp. B) Collection of laboratory *B. subtilis* strains was evaluated for presence of unique phi3T sequence using oAD38/oAD39 (expected product size – 1121 bp) (I); Strain 168 New Castle, which was positive for phi3T, was evaluated with additional primer sets to check for the presence of intact *kamA* and prophage integration into *kamA* gene (II); Difference between phi3T (KY030782.1) and phi3Ts was confirmed by primer pair s (oAD51/oAD52, expected product size for phi3Ts – 1557 bp; for phi3T and SPβ – 2097 bp). Similar to evolved strains, the 168 New Castle also appeared to carry phi3Ts (III).

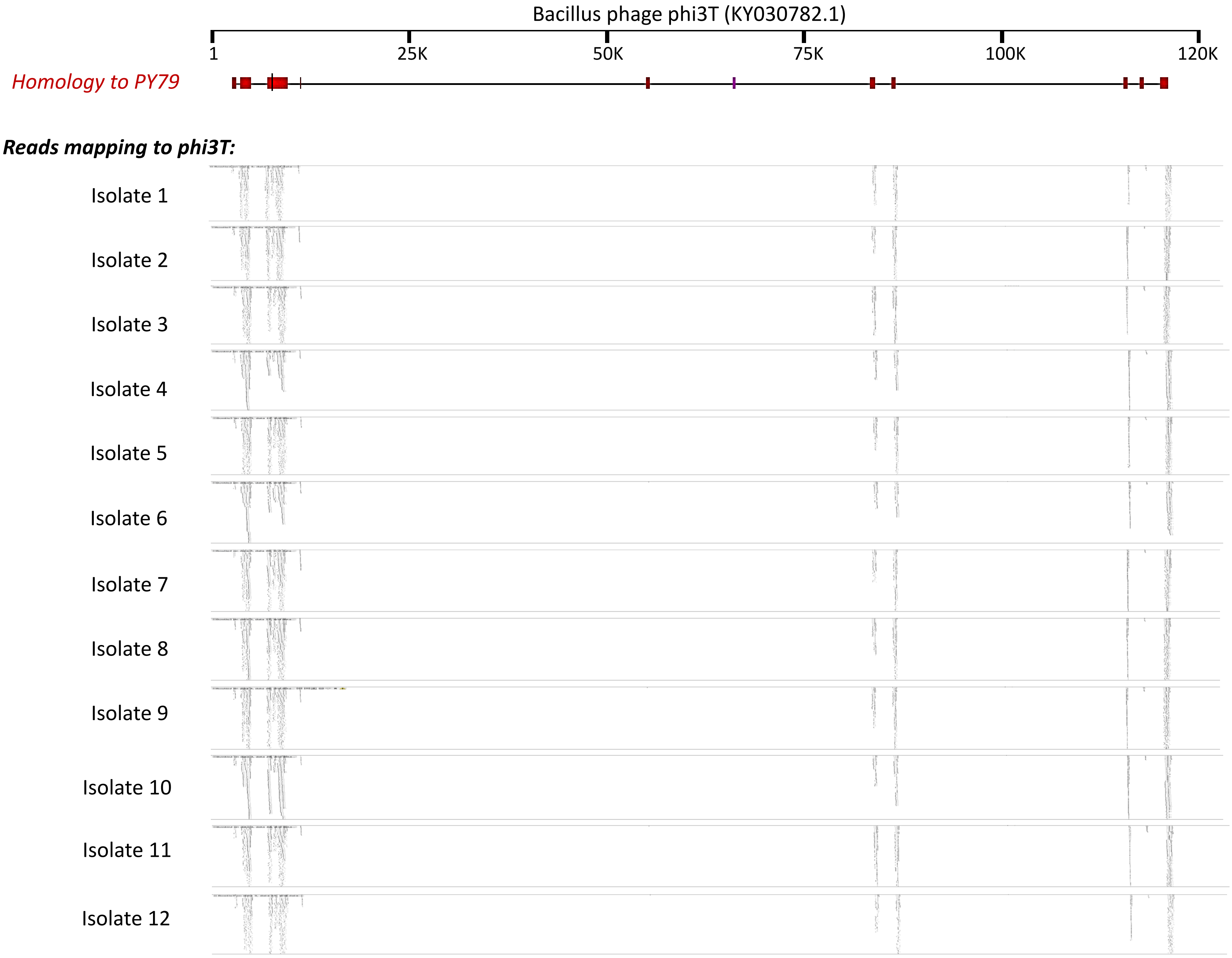


Supplementary Figure 5. Detection of phi3T DNA fragments in *B. subtilis* PY79 evolved under sporulation selection regime by another research group. Raw Illumina reads from the sequencing of 12 evolved isolates were mapped onto DNA of phi3T (KY030782.1) using Bowtie2 package in Galaxy platform (https://cpt.tamu.edu/galaxy-pub) and visualized using Trackster tool. Only reads, overlapping with high phi3T-PY79 homology regions were detected.

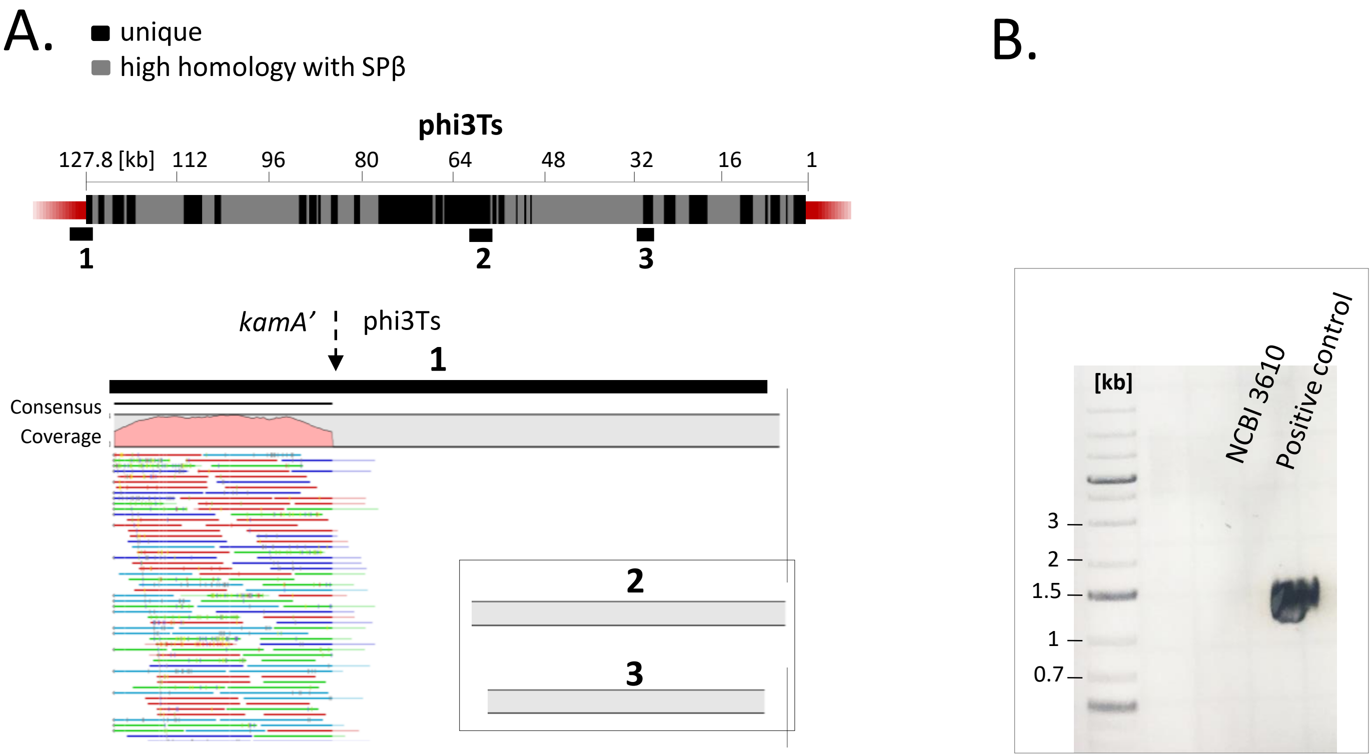


Supplementary Figure 6. Testing of WT NCBI 3610 for presence of phi3Ts. A) Detection of phi3Ts in WT NCBI 3610 strain through mapping of raw sequencing reads (just like performed for *B. subtilis* 168 ancestor, Fig 3). B) Detection of phi3Ts-specific PCR product using oAD38/oAD39 in NCBI 3610.

A.

| transfer nr => | I | II | III | IV | V | VI | VI | VIII |
|---|---|---|---|---|---|---|---|---|
| Strain (treatment) | | | | | | | | |
| | - | - | - | - | - | - | - | - |
| **WT (nt)** | - | - | - | - | - | - | - | - |
| | - | - | - | - | - | - | - | - |
| | - | - | +/- | + | - | - | - | - |
| **WT (heat)** | - | - | +/- | + | - | - | - | - |
| | - | - | +/- | + | - | - | - | + |
| | - | - | +/- | + | + | + | +/- | + |
| **WT (NaOH)** | - | - | +/- | + | - | + | + | + |
| | - | - | - | + | - | + | + | + |
| | - | - | - | - | - | - | + | - |
| **ΔSPβ (heat)** | - | - | - | - | + | - | + | - |
| | - | - | - | - | - | + | - | - |

B.

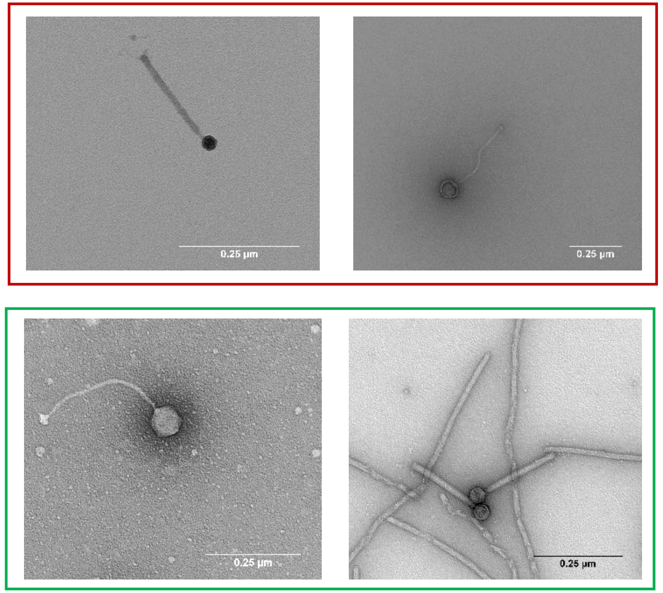


Supplementary Figure 7. Phage release by WT NCBI 3610 as a consequence of sporulation selection regime. A) WT NCBI 3610 was subjected to serial transfer experiment using three alternative treatments: nt – non-treated, where late-stationary phase cells were transferred into the fresh medium; heat – where cells were heat-treated prior the transfer selecting for dormant spores; NaOH – where cells were treated with 3 M NaOH, also selecting for dormant spores. In addition, the ΔSPβ strain was serially transferred with heat treatment, to evaluate if presence to this prophage is necessary for increase in lytic activity and occurrence of phage particles in the medium. B) Transmission electron microscopy images of phage precipitates obtained from WT (NaOH treatment) and ΔSPβ (heat treatment), confirming the presence of two types of phage particles in the medium (just like previously shown for *B. subtilis* 168, evolved under sporulation selection regime).

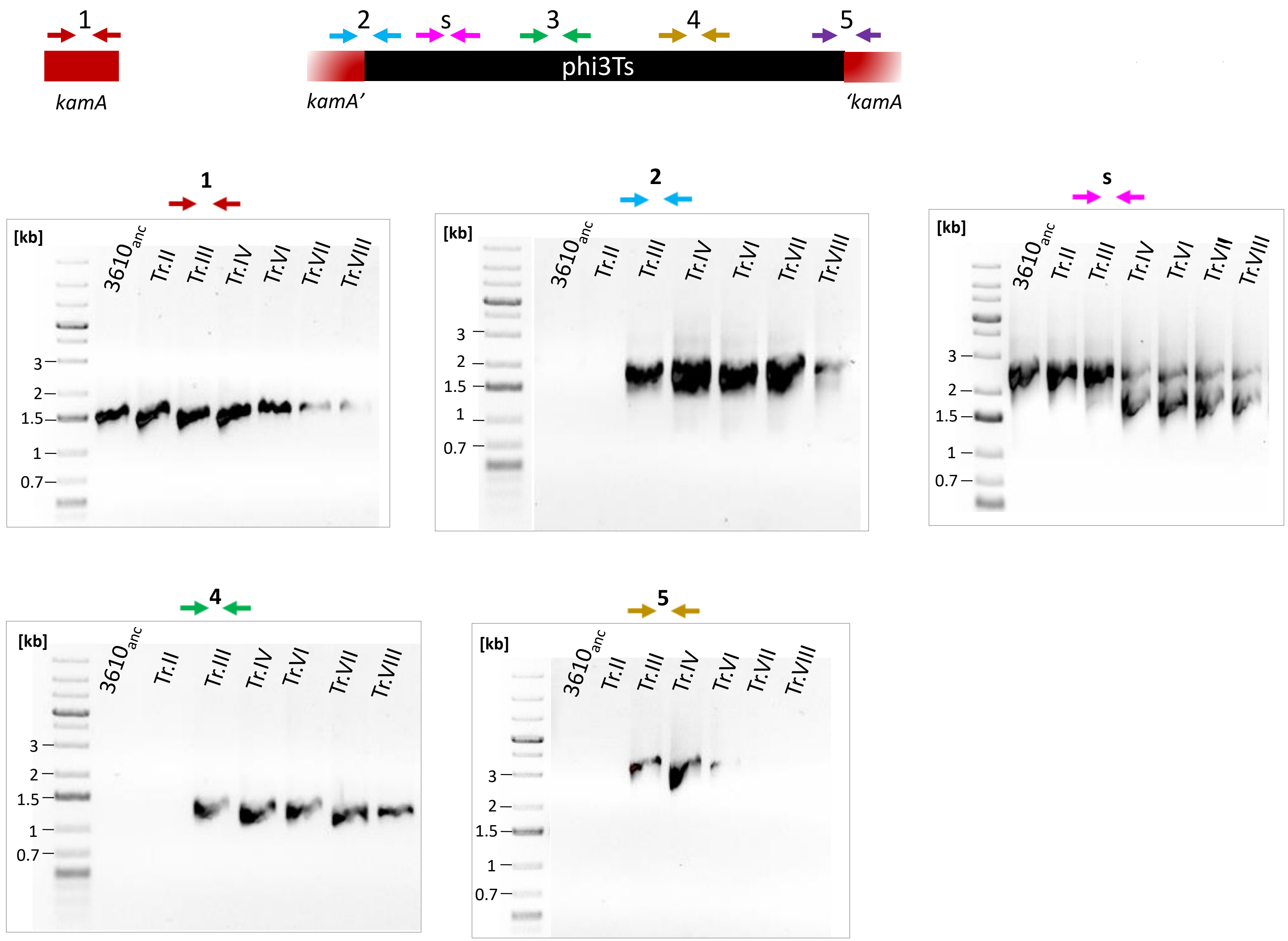


Supplementary Figure 8. Emergence of phi3Ts-specific PCR products upon sporulation selection regime. DNA was isolated from the ancestor and populations of cells from subsequent transfers, where NaOH spore selection regime was applied. Primer pairs used and expected molecular mass of the PCR product were as follows: 1 – oAD36/oAD37, 1366 bp; 2 oAD42/oAD43, 1819 bp; 3-oAD38/oAD39, 1121 bp; 4- oAD40/oAD41, 3033 bp; 5- oAD44/oAD45, 901 bp. B) Collection of laboratory *B. subtilis* strains was evaluated for presence of unique phi3T sequence using oAD38/oAD39 (expected product size – 1121 bp); s - oAD51/oAD52, expected product size for phi3Ts – 1557 bp; for phi3T and SPβ – 2097 bp.

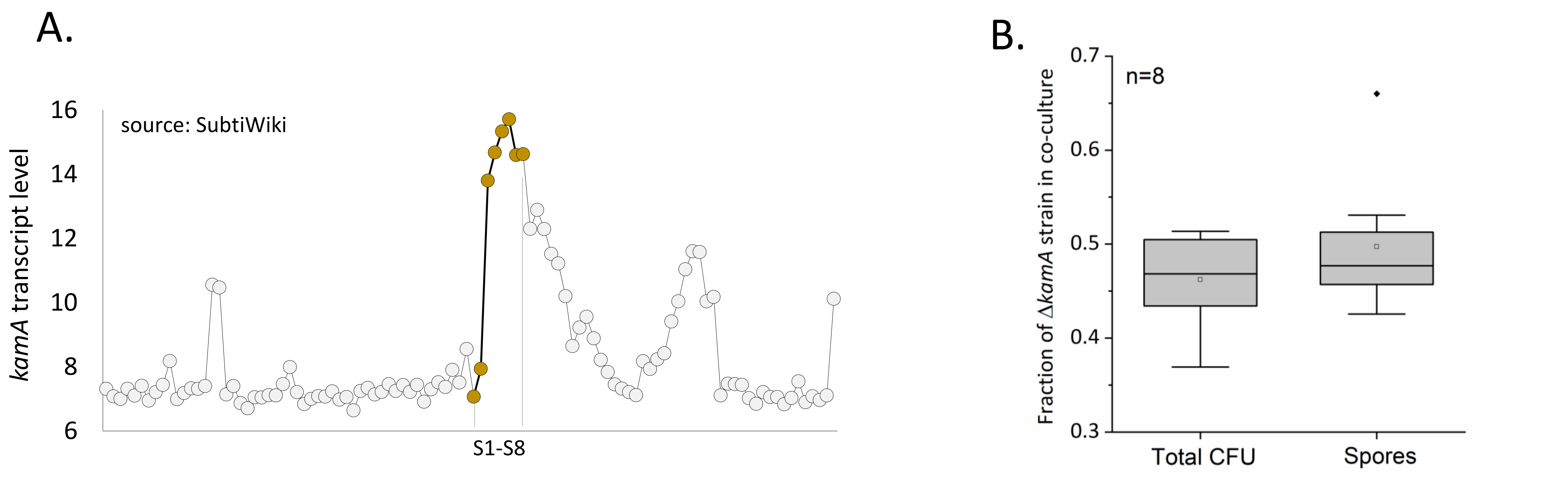


Supplementary Figure 9. Role of *kamA* in growth and sporulation. A) Expression profile of *kamA* gene, obtained from SubtiWiki website (http://subtiwiki.uni-goettingen.de), downloaded on 19.02.2020. Each for represents different experimental conditions. Dots highlighted in gold represent subsequent hours (from 1 to 8h) after entry into sporulation. B) Competition between WT and Δ*kamA* strain, starting from 1:1. After 48h the ratio between total populations as well as spore populations of WT and Δ*kamA*, remained not significantly different from 1:1 ($P<0.066$ and $P<0.92$, respectively). Boxes represent Q1–Q3 (quartiles), lines represent the median, and bars span from max to min. Dots represent outlier data points.

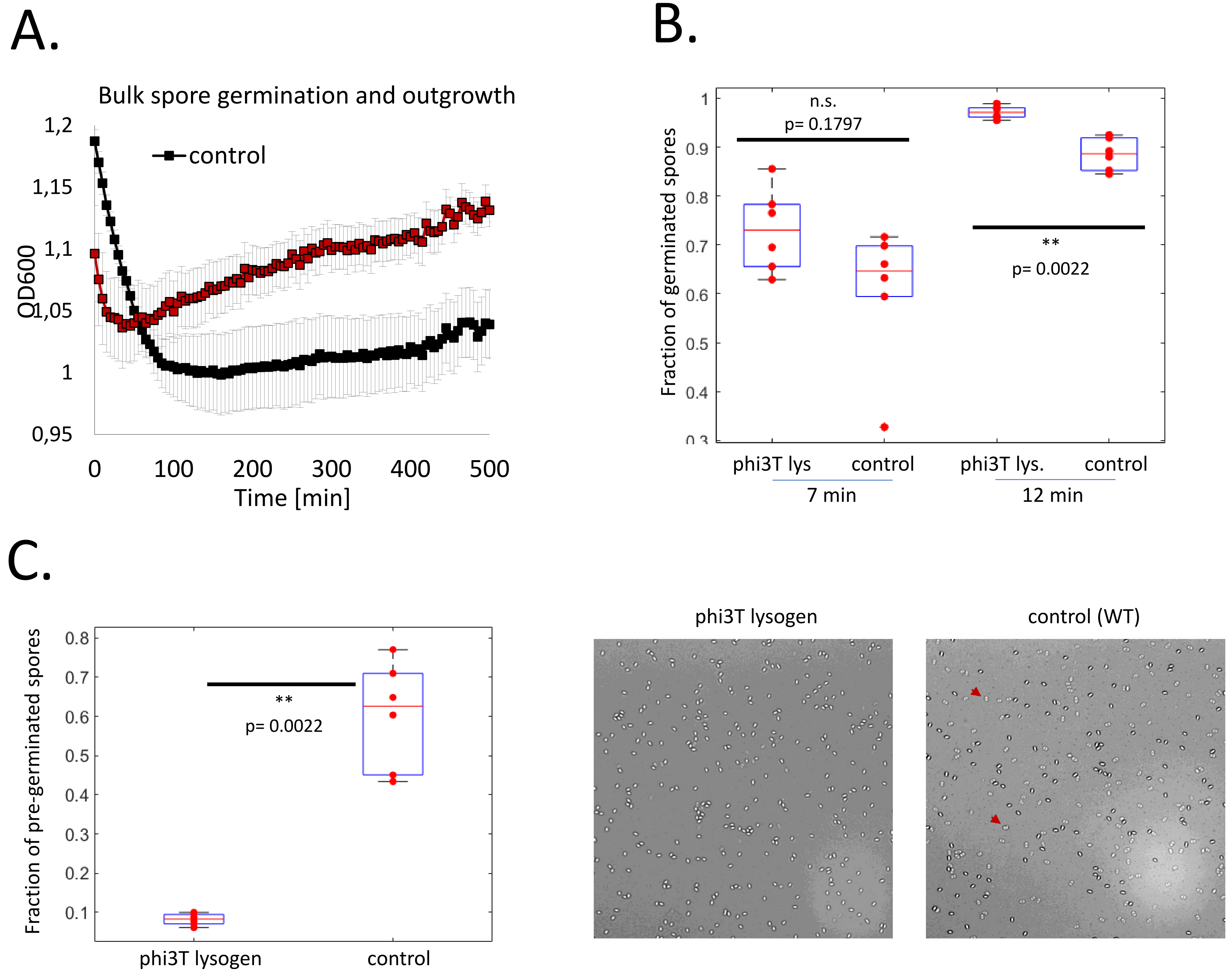


Supplementary Figure 10. Comparison of spore revival traits of WT (NCBI 3610) and phi3T lysogen. A) Bulk spore germination dynamics and outgrowth in liquid germination solution, monitored through changes in optical density. Germination results in a drop of OD which is followed by rising OD due to spore outgrowth and initiation of vegetative growth. Faster outgrowth of phi3T lysogen after germination is evident from slope differences between control (P<6.1□10-5) and the phi3T lysogen (P<1□10-4), but not statistically significant (P<0.07). B) Percentage of germinated spores upon induction of spores on agarose pads with L-alanine (see Methods for details). Dots represent data from individual images taken at 7 minutes and 12 minutes, respectively containing n > 60 spores from two technical replicates each (in total 766 and 1588 observed spores for WT and the phi3T lysogen, respectively). Percentage of induced germination was computed by excluding pregerminated spores. C) Left: Percent of phase-bright pre-germinated spores in uninduced sample of the WT and phi3T lysogen. Right: Representative brightfield microscopy images of spores derived from both strains. Red arrows point towards the pre-germinated spores.

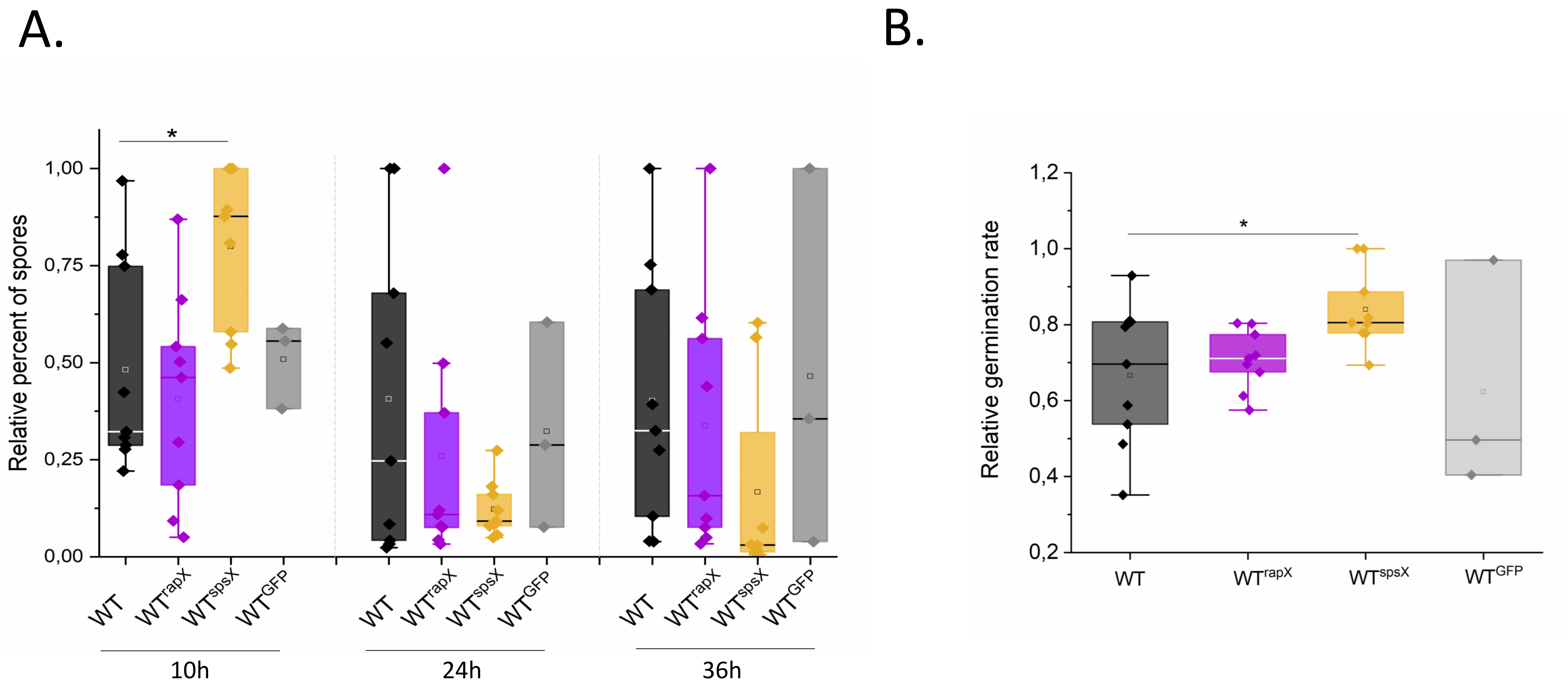


Supplementary Figure 11. Comparison of spore frequency and spore revival traits of WT (NCBI 3610), WT$^{rapX}$, WT$^{spsX}$ and WT$^{GFP}$ (as a control) and phi3T lysogen. A) Percent of spores was assessed after 10, 24 and 36h. Percent of spore values were collected from 3 independent experiments and represented in relative units (divided by maximal value within experiment in given timepoint). Strain WT$^{spsX}$ shows significantly higher percent of early spores compared to WT ($p<0.013$, Student's t test). B) Bulk spore germination dynamics and outgrowth in liquid germination solution was monitored through changes in optical density. Germination results in a drop of OD which is followed by rising OD due to spore outgrowth and initiation of vegetative growth. Slope values of declining ODs were collected from 3 independent experiments and represented in relative units (divided by maximal value within experiment) were collected, Germination of WT$^{spsX}$ is significantly faster compared to WT ($p<0.027$, Student's t test).

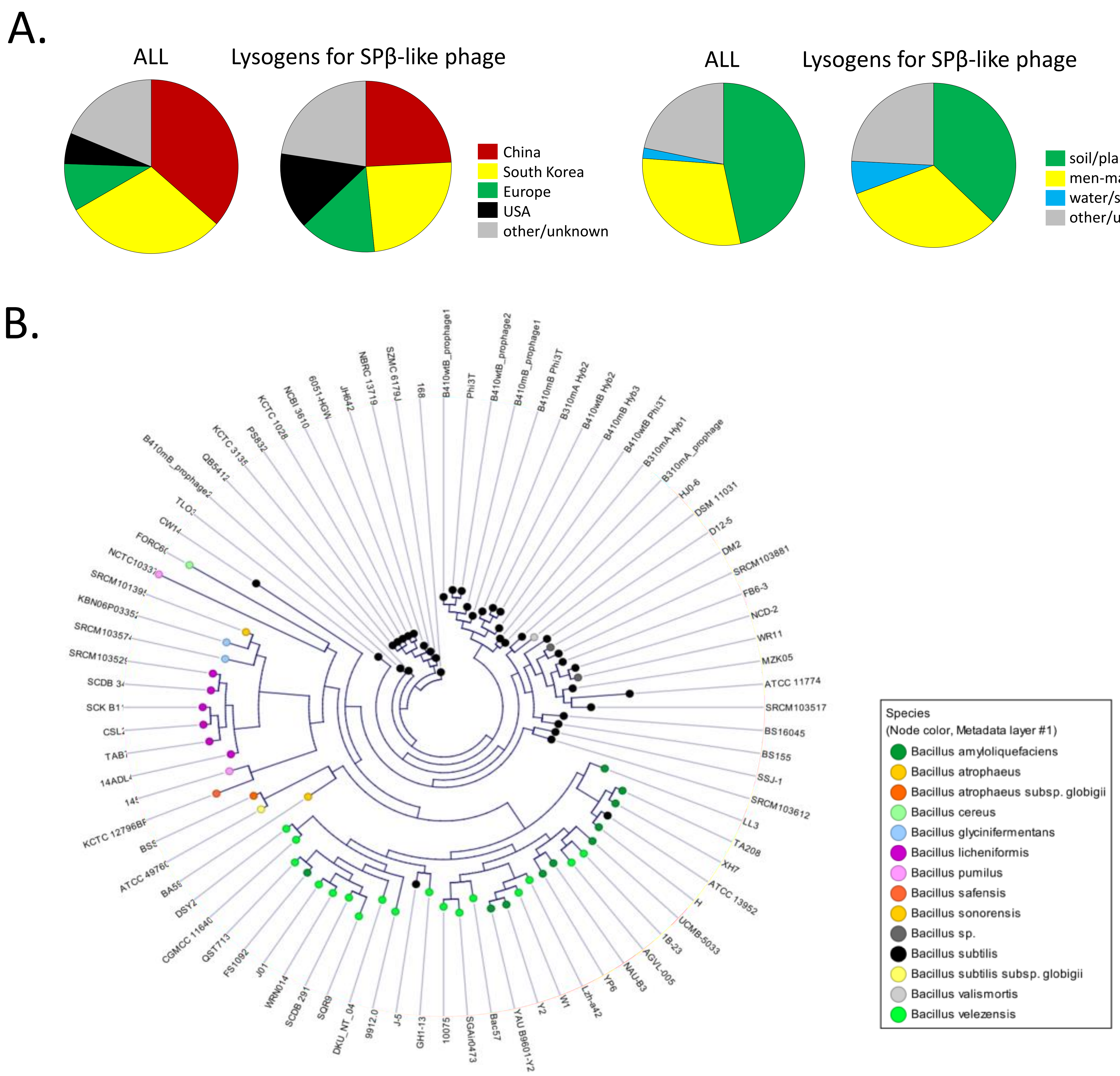


Supplementary Figure 12. Natural *Bacillus* sp. lysogens that carry a large prophage, integrated close to replication terminus (like SPβ). A) Comparison of geographical distribution of all *Bacillus* sp. strain included in the prophage elements analysis and strains that are lysogenic for large prophage. B) Comparison of isolation source of all *Bacillus* sp. strain included in the prophage elements analysis and strains that are lysogenic for large prophage. B) Phylogenetic tree of large *Bacillus* sp. prophages integrated close to replication terminus. Prophage elements cluster according to species.

**Supplementary Table 1 |** List of extrachromosomal phage DNA (epDNA) and hybrid phages, with indicated fragments of homology to SPβ region and phi3Ts, along with homologous (1-16) and non-homologous (N) junctions.

| **epDNA** | | **JUNCTION I** | | **JUNCTION II** | | **JUNCTION III** | | **JUNCTION IV** | |
|---|---|---|---|---|---|---|---|---|---|
| C15 (66.1 kB) | *cwlP/* 2250088-2264649 *blyA* (14561 bp) | **1 (5852 bp)** | *un/* 17903 -start (17903 bp) | ***attR*** | *kamA/* 2139866-2179307 */yorL* (39441 bp) | | | | |
| C22 (37.5 kB) | *cwlP/* 23720-start-end-112345 */reductase* (28623 bp) | **2 (1633 bp)** | *yotI* 2153661-2264204 *yosQ* (10534 bp) | | | | | | |
| C24 (53.3 kB) | *un/*20521-start-end-122932 */un* (25434 bp) | **3 (1637 bp)** | *yotI* 2153648 – 2172784 *mtbP* (19136 bp) | **4 (2467 bp)** | *un* 107106-94240 */yorI* (12866 bp) | | | | |
| C5 (27.3 kB) | *uvrX/* 9762-start-end- 122932 */un* (14665 bp) | **3 (1637 bp)** | *yotI* 2153648 – 2164642 *yosL* (10994 bp) | **5 (6851 bp)** | *un/* 118762-108609 */un* (10135 bp) | | | | |
| C25 (31.8 kB) | *yojW/* 2158201 - 2170033 *yosA* (11832 bp) | **N1** | *un/* 107382-87412 */un* (19970 bp) | | | | | | |
| C19 (56.4 kB) | *yorQ* 102540-84628 */un* (17912 bp) | **6 (1851 bp)** | *yoqW/* 2191852-2217032 *yopA* (25180 bp) | **7 (30 bp)** | *un/* 62446-47247 */yonJ* (15199 bp) | | | | |
| C18 (31.6 kB) | *yosT/* 117689-104631 *mtbP* (13058 bp) | **8 (2475 bp)** | *yorY* 2170317 to 2174332 *yorP* (4015 bp) | **9 (1415 bp)** | *yorS/* 103574- 85122 /ligase (18452 bp) | | | | |
| C14 (31.4 kB) | *un/* 127033-111907 */reductase* (15126 bp) | **10 (6855 bp)** | *yosV* 2157793-2180996 */yorK* (23203 bp) | | | | | | |
| C3 (46.1 kB) | *kamA/* 2139866-2164642 *yosQ* (24776 bp) | **5 (2475 bp)** | *un/* 118762- 104631 *mtbP* (14131 bp) | **8 (6849 bp)** | *yorY* 2170317-2186861 *yorE* (16544 bp) | | | | |
| C0 (26.2 kB) | *kamB/* 2141507 -2139866 */kamA* (1641 bp) | ***attR*** | start-24562 */cwlP* (24562 bp) | | | | | | |
| C1 (24.9 kB) | *un/* 91767-84628 *un* (7139 bp) | **6 (1895 bp)** | *yoqW/* 2191652-2202762 *yopW* (11110 bp) | **11 (3225 bp)** | *yoqD* 79566-67753 */un* (11813 bp) | | | | |
| C6 (36.7) kB | *un*/11510-start-end-120530 *thyA* (18805 bp) | **12 (400 bp)** | *sspC* 2156750-2164642 *yosQ* (7892 bp) | **5 6860 bp** | *un/* 118762-99485 */yorL* (19277 bp) | | | | |
| C410wtB (31.8 kB) | *un/* 55472-29334 *yozP* (26138 bp) | **13 (19685 bp)** | *yonK* 2226855-2252386 */cwlP* (25534 bp) | | | | | | |
| **Phages** | | | | | | | | | |
| Hyb1$^{\text{phi3Ts-SP}\beta}$ | *kamA/* 2138666-2139870 */yodN* (1204 bp) | ***attL*** | end-29326 *yozP* (98509 bp) | **14 (19758 bp)** | *yonK* 2226851- 2278535 *yoyK* (51684 bp) | | | | |
| Hyb2$^{\text{phi3Ts-SP}\beta}$ | end-*111907 /reductase* (15928 bp) | **10 (6855 bp)** | *yosV* 2157793-2172793 *mtbP* (15000 bp) | **4 (2467 bp)** | *un* 107106-start (107106 bp) | | | | |
| Hyb3$^{\text{phi3Ts-SP}\beta}$ | end-*111907 /reductase* (15928 bp) | **10 (6855 bp)** | *yosV* 2157793-2172793 *mtbP* (15000 bp) | **4 (2467 bp)** | *un 107106-54990 /un* (52116 bp) | **15 (518 bp)** | *yonP* 2221684-2246610 *yozP* (24926 bp) | **16 (19760 bp)** | start-49086 *yonK* (49086 bp) |

Fragments of homology to SPβ region (or region adjacent to SPβ) in *B. subtilis* (NC_000964.3), were labelled in pink, while fragments of homology to phi3Ts (MT366945.1) were labelled in grey. Numeric range indicates the corresponding regions of homology *in B. subtilis* or phi3Ts. The first and last gene contained within the homology fragment was indicated before and after the numeric range, respectively. Dash sign (/) indicates that the gene is interrupted, while lack of dash sign indicates that the fragment starts or ends in the intergenic region. Numbers in brackets indicate sizes of each homologous fragment or junction. Unique junctions are numbered from 1-16/N1.

**Supplementary Table 2 | Predicted function of SPβ and phi3Ts gene (RAST annotation)**

| | |
|---|---|
| ***Present in SPβ and phi3Ts*** | |
| *yotM* | putative recombinase |
| *sspC* | small acid-soluble spore protein |
| *yosT* | transcriptional regulator, AraC family |
| *nrdIB* | ribonucleotide reductase of class Ib |
| *yorR* | putative nucleotide kinase |
| *recJ* | single-stranded DNA-specific exonuclease |
| *yorI* | phage-associated DNA helicase |
| *ligB* | bacteriophage SPβ DNA ligase |
| *yoqD* | putative nucleotide kinase |
| *yonN* | DNA binding protein HBsu |
| *yonF* | putative prophage terminase, ATPase subunit |
| *yonD* | phage virion protein |
| *yomS* | putative phage lytic exoenzyme |
| *yomM* | putative integrase |
| *cwlP* | N-acetylmoramoyl-L-alanine amidase |
| *uvrX* | putative UV-damage repair protein |
| *bsrG* | type I toxin-antitoxin system |
| *yokF* | chromosome-degrading nuclease |
| ***Unique for SPβ*** | |
| *yotN* | accessory protein for the excision of the SPβ prophage |
| *yosA* | putative type I toxin |
| *aimP* | arbitrum peptide |
| *aimR* | arbitrum transcriptional regulator |
| *yonR* | putative transcriptional regulator (Xre family) |
| *yomJ* | putative phage immunity protein |
| *bdbA* | bacteriophage thiol-disulfide-oxidoreductase |
| *sunA* | sublancin antibiotic precursor |
| *yoyK* | putative DNA wielding protein |
| *yokA* | site specific recombinase |
| ***Unique for phi3Ts*** | |
| *thyB* | thymidylate synthase |
| *spsX* | stationary phase survival protein |
| *aimP* | arbitrum peptide |

| | |
|---|---|
| *aimR* | arbitrum transcriptional regulator |
| *traIS3* | IS3 family transposase |
| *rapX* | putative aspartate phosphatase, sporulation regulator |
| *pinR* | site-specific recombinase |

**Supplementary Table 3 | The strains used in this study.**

| ***Strains used in experiments*** | | |
|---|---|---|
| Strain name | Genotype | Reference |
| B310mA | Δ*eps*-Δ*tasA* evolved under sporulation selection regime | [1] |
| B410mB | Δ*eps*-Δ*tasA* evolved under sporulation selection regime | [1] |
| B410wtB | 168 hymKATE $P_{tapA\text{-}yfp}$ | [1] |
| DK1042 | Naturally competent derivative of the undomesticated NCIB 3610 containing *comI*$^{Q12I}$ allele | [2] |
| DTUB200 | 3610 *comI*$^{Q12I}$ infected with Bacillus phage phi3T | this work |
| DTUB201 | 3610 *comI*$^{Q12I}$ SPβ*::ery* | this work |
| DTUB202 | 3610 *comI*$^{Q12I}$ *kamA::km* | this work |
| DTUB251 | 3610 *comI*$^{Q12I}$ *amyE::* $P_{hyperspank}$-*rapX* (Spec$^R$) | this work |
| DTUB254 | 3610 *comI*$^{Q12I}$ *amyE::* $P_{hyperspank}$-*spsX* (Spec$^R$) | this work |
| TB500 | 3610 *comI*$^{Q12I}$ *amyE::* $P_{hyperspank}$-GFP (Spec$^R$) | [3] |
| Δ6 | *trpC2*; ΔSPβ; sublancin 168-sensitive; Δ*skin*; ΔPBSX; Δprophage1; *pks*::Cm; Δprophage 3; Cm$^r$ | [4] |
| ***Strains used as phage or gDNA donors*** | | |
| CU1065 | (phi3T) attSPβ trpC2 | BGSC |
| BKK19690 | *kamA*::*km* trpC2 | BGSC |
| SPmini | SPβ*::ery* | [5] |

**Supplementary Table 4 | Phages used in this study.**

| Name | Description | Reference |
|---|---|---|
| phi3T | Bacillus phage isolated from CU1065 lysogen | [6] |
| phi3Ts | Bacillus phage isolated from B410mB lysogen | this work |
| Hyb1$^{phi3Ts\text{-}SP\beta}$ | Bacillus hybrid phage released by B310mA | this work |
| Hyb2$^{phi3Ts\text{-}SP\beta}$ | Bacillus hybrid phage released by B310mA and B410wtB | this work |
| Hyb3$^{phi3Ts\text{-}SP\beta}$ | Bacillus hybrid phage released by B410mB | this work |

**Supplementary Table 5 | Primers and plasmids used in this study.**

| ***Primers*** | | |
|---|---|---|
| Symbol | Experimental purpose | Sequence (5' to 3') |
| oAD36 | Verify the integrity of *kamA* gene (phi3Ts insertion side) | CCGCATTCAGTCTCTTTC |
| oAD37 | Verify the integrity of *kamA* gene (phi3Ts insertion side) | GGAAGGAGATCGAGTTAT GG |
| oAD38 | Confirm presence of phi3Ts unique fragment I (no homology to SPβ) | CTCTGTGGGCATCACTTC |
| oAD39 | Confirm presence of phi3Ts unique fragment I (no homology to SPβ) | CTGGTAGCTCAGCTAAAG |
| oAD40 | Confirm presence of phi3Ts unique fragment II (no homology to SPβ) | GGTTGAAGACGGACTGAA G |
| oAD41 | Confirm presence of phi3Ts unique fragment II (no homology to SPβ) | ACGGAGTCTGCGTAATG |
| oAD51 | Check for presence of unique *spsX* gene (only in phi3Ts) | AGAGCCGGTCAAAGGTAA AC |
| oAD52 | Check for presence of unique *spsX* gene (only in phi3Ts) | GCTTTGCTGCAACTGTTG |
| oAD47 | Gibson cloning to open pDR111 plasmid | TCGACTAAGCTTAATTGTT ATCC |
| oAD48 | Gibson cloning to open pDR111 plasmid | CATGCAAGCTAATTCGGTG G |
| oAD49 | Gibson cloning to insert *rapX* into pDR111 | GACCTCGTTTCCACCGAAT TAGCTTGCATGTTAAGTTT CATAAAGACAATCCCCTCT CTG |
| oAD50 | Gibson cloning to insert *rapX* into pDR111 | TGTGAGCGGATAACAATTA AGCTTAGTCGAAGGGCTTG TGTTGGAGCAGATG |
| oAD71 | Gibson cloning to insert *spsX* into pDR111 | GACCTCGTTTCCACCGAAT TAGCTTGCATGCCCTAATT AATAATTGAAACCGTTCCA TG |
| oAD72 | Gibson cloning to insert *spsX* into pDR111 | TGTGAGCGGATAACAATTA AGCTTAGTCGAGCTATGAT |

| | | |
|---|---|---|
| | | AATTTTAATCCCACTGGCA AC |
| ***Plasmids*** | | |
| Name | Host | Purpose |
| pDR111_rapX | *E. coli* MC1000 | Complementation of *B. subtilis* with phi3Ts putative regulator RapX |
| pDR111_sspX | *E. coli* MC1000 | Complementation of *B. subtilis* with phi3Ts putative regulator SpsX |
